# Identification of the Rigid Core for Aged Liquid Droplets of the TDP-43 Low Complexity Domain

**DOI:** 10.1101/2021.03.01.433427

**Authors:** Blake D. Fonda, Khaled M. Jami, Natalie R. Boulos, Dylan T. Murray

**Author notes:** **Corresponding Author:** Dylan T. Murray, Department of Chemistry, University of California, Davis, CA 95616.

## Abstract

The biomolecular condensation of proteins with low complexity sequences plays a functional role in RNA metabolism and a pathogenic role in neurodegenerative diseases. The formation of dynamic liquid droplets brings biomolecules together to achieve complex cellular functions. The rigidification of liquid droplets into β-strand-rich hydrogel structures composed of protein fibrils is thought to be purely pathological in nature. However, low complexity sequences often harbor multiple fibril-prone regions with delicately balanced functional and pathological interactions. Here, we investigate the maturation of liquid droplets formed by the low complexity domain of the TAR DNA-binding protein 43 (TDP-43). Solid state nuclear magnetic resonance measurements on the aged liquid droplets identify a structured core region distinct from the region thought to be most important for pathological fibril formation and aggregation. The results of this study show that multiple segments of this low complexity domain are prone to form fibrils, and that stabilization of β-strand-rich structure in one segment precludes the other region from adopting rigid fibril structure.

**Table of Contents Graphic:** 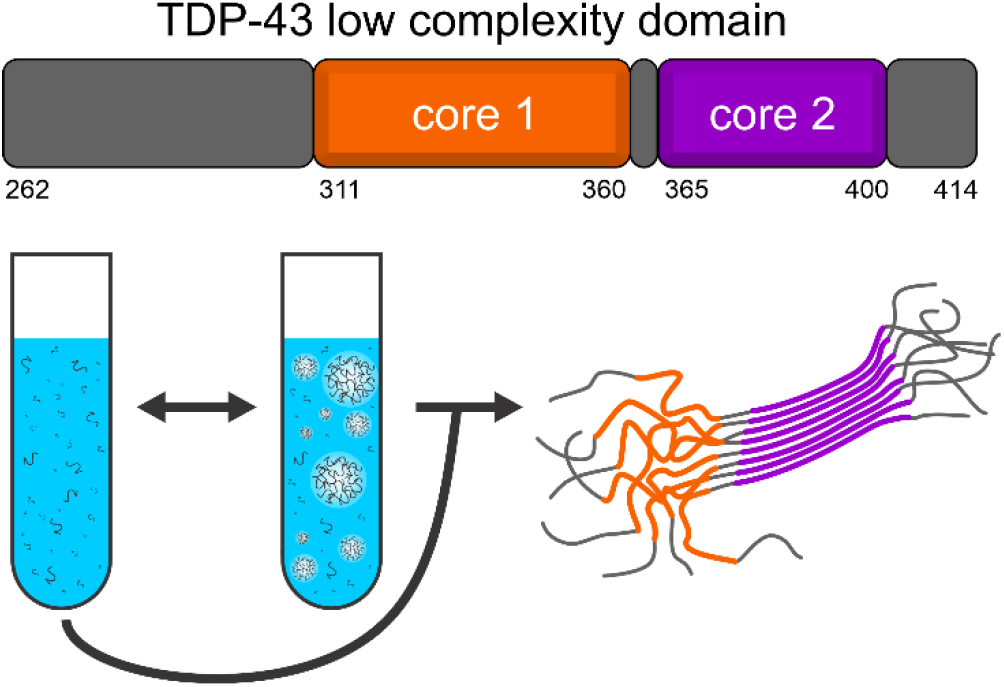

## Introduction

Biomolecular condensation is a broad term for chemical processes that living cells use to spatially and temporally localize molecules and chemical activity.^1^ Cellular functions such as transcriptional regulation, chromosome separation, neurotransmitter release, and the stress response are finely tuned implementations of these chemical processes.^2^ These processes rely on a variety of weak and transient multivalent interactions between the molecules involved.^3^ Due to its prevalence in biological systems, biomolecular condensation has developed into a highly active area of chemical research. Common themes regarding the underlying microscopic molecular interactions responsible for this behavior are emerging, yet significant questions remain unanswered for specific biological systems.^3,4^

Granular structures in living cells called membraneless organelles have macroscopic properties consistent with biomolecular condensations.^5^ The self-association of protein sequences with reduced amino acid complexity is paramount to the formation of granular structures involved in the processing and transport of RNA molecules.^6^ These intrinsically disordered low complexity (LC) protein domains are ubiquitous in the genomes of higher eukaryotes.^7^ *In vitro*, LC domains form biomolecular condensates with varying degrees of structure and molecular order that exhibit the same phenomenological behavior as the *in vivo* RNA granules.^8^ LC domains transition from a monomeric state characterized by intrinsic disorder to a highly dynamic self-associated state formed through a liquid-liquid phase separation process, commonly referred to as a liquid droplet. If the molecular interactions responsible for liquid droplet formation are left unregulated, the LC domains can lose their dynamics and transition to more rigid hydrogel assemblies.^1^

While liquid droplets are usually associated with functional activity, their conversion into more rigid hydrogels is thought to play a role in disease pathology. The fused in sarcoma (FUS) RNA-binding protein forms liquid droplets that age into hydrogels.^9^ This process occurs both in living cells and with purified proteins, and is sped up by mutations in the LC domain linked to amyotrophic lateral sclerosis (ALS).^9^ Solution nuclear magnetic resonance (NMR) measurements indicate that the LC domain inside the liquid droplets is primarily disordered^10^ and solid state NMR measurements show that the hydrogels are composed of LC domains in an amyloid-like cross-β fibril conformation.^11^ Disease mutations in LC domains disrupt native interactions and molecular motions, favoring the more rigid fibrillar hydrogel state, consistent with the buildup of pathological aggregates in patients harboring these mutations.^6^ For FUS and similar proteins, common motifs in LC domains facilitate the formation of both liquid droplets and hydrogels. For example, chemical footprinting experiments of the LC domain in the heterogeneous nuclear ribonucleoprotein A2 (hnRNPA2) show that a specific sub-region of the LC domain is protected from chemical modification in liquid droplets, hydrogels, and nuclear condensates, revealing that this segment participates in the interactions that stabilize these assemblies.^12^ Electrostatic repulsion caused by residue-specific phosphorylation in the same 60 residue region of the 214 amino acid FUS LC domain prevents assembly both into liquid droplets and hydrogels, showing that close intermolecular association of these 60 residues is important for the formation of the macroscopic structures.^11^ LC domains represent functional self-assembling systems that form liquid droplets characterized by significant molecular motion and disorder in a delicate balance with a propensity to form hydrogels composed of fibrils with limited molecular motion and significant order. Disruption of this balance results in pathological aggregates that are the hallmark of devastating neurodegenerative diseases.

The 414 residue RNA-binding protein TAR DNA binding protein 43 (TDP-43) is a primary component of RNA granules and has functions in transcription, translation, and splicing.^13^ TDP-43 forms pathological aggregates in neurodegenerative diseases such as ALS,^14,15^ frontotemporal dementia (FTD),^14,15^ and Alzheimer’s Disease.^16^ The full-length protein has four main domains, an N-terminal oligomerization domain, two RNA recognition motifs (RRMs), and a C-terminal LC domain (TDP-43-LC).^6^ The LC domain, residues 274–414, is necessary for the formation of both liquid droplet and hydrogel structures. The functional activities of TDP-43-LC are typically associated with the formation of liquid droplets, and the rigidification of these structures is associated with disease pathology.^13^

Almost all ALS and FTD related missense mutations lie in TDP-43-LC.^6^ Deletion of residues 311–360 of TDP-43-LC prevents the formation of aggregates in living cells.^17^ Residues 311–360 are also vital for the formation of both liquid droplets and hydrogels. Solution NMR chemical shift measurements indicate that this region transiently samples a helical conformation in monomeric TDP-43-LC.^18^ These experiments also show oligomeric species form in a concentration-dependent manner through transient contacts in the helix-prone region.^18^ Mutations in the helix-prone region disrupt liquid droplet formation for TDP-43-LC, but aging and rigidification of the mutant liquid droplet samples preclude direct observation of the helix in this state via solution NMR.^18^ An additional interaction site near the C-terminal region of the LC domain in residues 382–385 transiently contacts the helix-prone region and is thought to have a role in the formation of the liquid droplets.^18^ Another study shows TDP-43-LC forms fibrils with a characteristic diffraction pattern and Thioflavin T (ThT) binding properties indicative of cross-β fibril formation which matches the binding properties of neuronal inclusions obtained from patient tissues in autopsy.^19^ Experiments on residues 311–360 of TDP-43-LC indicate that in the absence of the flanking regions, the LC domain converts to a β-strand conformation as it transitions from liquid droplet to fibril morphology.^20^ Solid state NMR measurements indicate residues 311–360 converge on a single conformation in fibril form^20^, while cyro-electron microscopy (cryo-EM) structures of this fragment reveal this segment forms polymorphic fibrils.^21^

Structural polymorphism is a common trait of pathogenic fibril forming proteins. Examples of these proteins include the β-amyloid, tau, and α-synuclein proteins.^22^ The propensity of residues 311–360 in TDP-43-LC to adopt multiple conformations^21^ may be indicative of a purely pathological process. However, the footprinting studies on hnRNPA2^12^ and the phosphorylation experiments on FUS^11^ suggest that the conversion of TDP-43-LC into fibrils may be the result of functional interactions left unregulated in the test tube. The full-length LC domains of the FUS and hnRNPA2 proteins do not exhibit polymorphism and condense into a single conformation using the same segments that are most important for liquid droplet formation.^11,23^ For FUS, removal of the monomorphic core forming segment allows other regions of the LC domain to adopt stable fibril-like conformations.^7,24,25^ The presence of more than one stable core in a LC domain could determine the balance between functional and pathological assembly processes. For example, the propensity to form structure in a non-pathogenic core segment could prevent detrimental aggregation from pathogenic regions of a LC domain. Disease mutations disrupting this balance could then lead to pathogenic aggregation.

Does the complete LC domain of TDP-43 behave in the same manner? Does a non-pathogenic core in the TDP-43-LC prevent pathological aggregation? In this paper, we characterize the structures of TDP-43-LC fibrils formed through both aging of liquid droplets and a ‘flash diluted’ protocol that circumvents a prolonged dynamic phase. These measurements provide insight into the balance of interactions present across the complete LC domain. Our results are in stark contrast to existing studies of TDP-43-LC domain aggregation and provide a basis for understanding functional to pathogenic transitions for TDP-43-LC.

## Experimental Section

### Protein expression and purification

His-tagged TDP-43-LC, in a pHIS-parallel plasmid containing residues 262–414 of human TDP-43 was expressed in BL21(DE3) cells. All bacterial growth was conducted at 37 °C with shaking at 220 RPM. Bacteria cells were grown in 4 l of Luria broth media to an OD_600_ of ~1.0. Cells were harvested at 6,000 g for 10 min and then transferred to 1 l of M9 minimal media containing U-^13^C_6_ glucose and ^15^N ammonium chloride for 30 min. Protein expression was induced by adding isopropyl-β-D-thiogalactoside (IPTG) to 0.5 mM. Cells were grown for an additional 3 h, harvested at 6,000 g for 10 min, flash frozen and stored at −75 °C for approximately 2 days before purification.

The cell pellet was thawed over ice and then resuspended in a solution containing 6 M guanidinium-HCl, 1% v/v Triton X-100, 500 mM sodium chloride, 50 mM Tris-HCl pH 7.5 via pipetting. Pierce Protease Inhibitor tablets were added according to the manufacturer’s specifications along with 0.25 mg/ml hen egg white lysozyme powder. The pellet was then sonicated in an ice bath using a Branson SFX-250 Sonifier with a 1/8-in microtip in pulsed mode with 0.3 s on and 3 s off, and 30% power for 20 min total ‘on’ time. The lysed cells were then centrifuged at 75,600 g for 30 min at 4 °C to remove non-soluble cell debris.

Protein purification was performed on a Bio-Rad NGC Discover 10 chromatography system. The supernatant was loaded onto a 5 ml Bio-Scale™ Mini Nuvia™ IMAC column equilibration in 6 M urea, 20 mM HEPES pH 7.5, and 500 mM NaCl. The column was washed with 6 M urea, 20 mM 4-(2-hydroxyethyl)-1-piperazineethanesulfonic acid (HEPES) pH 7.5, 500 mM NaCl, and 20 mM imidazole until the 280 nm absorbance returned to baseline. Protein was eluted from the column with 6 M urea, 20 mM HEPES pH 7.5, 500 mM sodium chloride, and 200 mM imidazole. SDS-PAGE analysis shows the sample purity is at least 95%.

### Liquid Droplet and Fibril Preparation

‘Flash diluted’ TDP-43-LC was prepared by diluting purified protein to a concentration of 3.4 mg/ml in a 3.5 ml volume in elution buffer (6 M Urea 500 mM NaCl, 200 mM Imidazole, 20 mM HEPES pH 7.5). The solution was transferred to a Millipore Amicon Ultra-15, 3 kDa MWCO, centrifugal filter and the buffer was exchanged into fibril buffer (20 mM sodium phosphate pH 7.4 and 200 mM sodium chloride) by centrifuging to a volume of 1 ml, resuspending to a volume of 4.5 ml with fibril buffer, centrifuging again to a volume of 0.8 ml, and resuspending to a volume of 4.3 ml. Approximately 4% of original elution buffer remained in the sample after buffer exchange. A cloudy solution containing white precipitate-like material was immediately observed. The solution was then homogenized via rotation for 4 d at room temperature (~20 °C). The fibril solution was then diluted two-fold with fibril buffer and sonicated with a Branson SFX-250 sonifier with a 1/8-in micro tip in pulsed mode at 15 % power, 1s “on”, 1s “off”, for a total time of 60s. The fibrils were again homogenized via gentle rotation for 12 d. To obtain sufficient material for solid state NMR measurements, a second batch of fibrils was prepared using the same procedure as the first with the following modifications: the centrifugal filter was a Millipore Amicon Ultra-4 3 kDa MWCO centrifugal filter, the initial homogenization step was 3 d, and the second, post-sonification, homogenization step was 5 d. Both samples were combined and packed into a solid state NMR rotor for measurements.

Liquid droplet TDP-43-LC was prepared by diluting purified protein to 2 mg/ml in elution buffer. The protein solution was loaded into 3.5 kDa MWCO dialysis tubing (Spectra Por 3 regenerated cellulose, flat width 29 mm) and dialyzed against fibril buffer for 52 h. The sample was harvested and then homogenized via gentle rotation for 92 h at room temperature.

### Electron Microscopy

TDP-43-LC samples were imaged using Ted Pella ultrathin carbon films (< 3 nm) on 400 mesh lacy carbon copper grids. Grids were glow discharged before adding 5 μl of undiluted protein solution and incubating for 2 min. The grid was then washed twice with 10 μl of ultra-pure water for ~10 s and then stained with 5 μl of a 3% (w/v) uranyl acetate solution for 10 s. After each solution was applied to the grid, bulk liquid was wicked away using a laboratory tissue prior to adding the next solution. The grid was air-dried prior to imaging on a JEOL-1230 electron microscope equipped with a 2k × 2k Tietz CCD camera operating at 100 keV.

### Liquid Droplet Imaging

TDP-43-LC liquid droplets were imaged using an Olympus BX51 light microscope with differential interference contrast (DIC) filters, a 40X objective lens, and a Diagnostics Instruments RT Slider camera with 6 megapixel sampling. For each series of images recorded, 3 μl of the TDP-43-LC solution was removed from the dialysis tubing and placed directly on standard untreated microscope glass (Fisher Scientific).

### Denaturant Quantification

Urea removal during dialysis was quantified by dialyzing of 8 ml of elution buffer against 1.5 l of fibril buffer in 3.5 MWCO kDa dialysis tubing (Spectra Por 3 regenerated cellulose, flat width 29 mm.) For each time point, 540 μl of the sample was removed from the dialysis bag, 60 μl of D_2_O was added, and a ^1^H 1D spectrum with water suppression was recorded using a 9.4 T Bruker Avance IIIHD Nanobay spectrometer and the WATERSUP pulse program. The percent change in urea concentration was determined by integrating the peak corresponding to urea protons (5.2 to 6.2 ppm) and dividing by the integral from the spectrum of undiluted elution buffer. Uncertainties in the integrals are less than 2.2% of the total area based on the RMS noise in individual 1D ^1^H spectrum.

### Fluorescence Measurements

Stock ThT solution, >500 μM, was prepared by dissolving solid ThT (Acros Organics) in ultrapure water (stirred for 1 h under ambient conditions). The stock solution was sterile filtered through a 0.22 μm PES membrane (Millipore). Working solution was prepared by diluting to 40 μM in Fibril buffer. Concentration of the diluted solutions were confirmed by absorbance at 412 nm using an extinction coefficient ε_412_ of 36,000 M^−1^cm^−1^ (Eppendorf BioSpectrometer Kinetic). All dye solutions were encased in aluminum foil and stored at 4 °C for up to 7 d. For ThT measurements, samples were prepared by adding 50 μl ThT working solution (40 μM ThT in Fibril Buffer) to 50 μl of protein sample. For intrinsic Trp measurements, the samples were loaded directly into the cuvette without dilution.

Both intrinsic Trp and ThT fluorescence measurements were recorded on an Agilent Cary Eclipse fluorescence spectrophotometer and a 45 μl 10 x 10 mm quartz cuvette (Starna). The excitation and emission slit widths were 5 nm, the scan rate was 600 nm/min, the averaging time was 0.1 s, and the data interval was 1.0 nm. Intrinsic fluorescence spectra were acquired between 300 and 500 nm, with excitation at 280 nm. ThT spectra were acquired between 455 and 600 nm, excitation at 440 nm. For the ThT data presented in Figure 2, the background fluorescence of the ThT solution without protein was subtracted from each spectrum.

### Solid State NMR Data Collection

8–10 mg of TDP-43-LC from the aged droplets or the flash-diluted preparation were packed into thin-walled 3.2 mm pencil style zirconia rotors (Revolution NMR) by centrifugation at 25,000 g for 1 h. 1–2 mm thick PTFE disks were used to center the sample in the rotor. NMR experiments were performed on an 18.8 T Bruker NMR spectrometer using an Avance III console and a BlackFox probe. MAS rates were 13.0 kHz for all experiments and cooling air was set so the sample temperature was ~15–20 °C. SPINAL64 decoupling^26^ was used for all direct and indirect data acquisition periods and continuous wave decoupling was used during specific CP (SCP)^27^. Indirect data was recorded using ‘States-TPPI’ mode. ^1^H T_1ρ_ values were determined using a 62.5 kHz ^1^H irradiation period of variable length before the CP step in a standard ^1^H-^13^C CP experiment. ^15^N T_2_ values were determined using a variable length echo delay after the ^1^H-^15^N CP step in a CP-NCA experiment. Relaxation decay curves were analyzed in the Topspin software (Bruker) and fit to exponential decay functions. Data were processed in nmrPipe^28^ and visualized in Sparky.^29^ All data collection and processing details for the NMR experiments presented in this paper are listed in Supplemental Table S6. The difference spectra were calculated by interpolating the aged liquid droplet spectra onto the same frequency grids as the flash diluted spectra using nmrPipe.^28^ The aged liquid droplet spectra were multiplied by a factor of 2.0 before subtracting the flash diluted spectra. The value of 2.0 was chosen to make the difference in the sharp signals between the spectra being compared negligible.

### Solid State NMR Signal Assignment and Torsion Angle Prediction

The reduced amino acid types present in TDP-43-LC (i.e. its low complexity character) make manual sequence specific assignment of the observed NMR chemical shifts challenging. To ensure an unambiguous placement of the observed signals in the TDP-43-LC primary sequence, signal assignments were determined using the MCASSIGN algorithm.^30^ Signal tables were generated from the 3D NCACX, NCOCX, and CANCO spectra of the flash diluted TDP-43-LC sample and are presented in Supplemental Tables S3, S4, and S5. Uncertainties in signal positions were determined based on resonance intensity and linewidth from the spectra, typically ranging from 0.3 to 0.4 ppm. For a subset of CO peaks in the NCACX spectrum larger uncertainties were assigned due to relatively weak signals or poor resolution of multiple signals. Several signals given a multiplicity of 2 in the NCACX and CANCO signal tables based on signal intensity and the results of preliminary MCASSIGN runs.

The MCASSIGN algorithm was run using the amino acid sequence for TPD-43-LC (residues 262-414) with the ‘good’, ‘bad’, ‘edge’, and ‘used’ weights ranging from (0, 60), (10, 60), (0, 8), and (0, 2), respectively, for 200 steps with 10^8^ iterations per step. 50 independent runs of MCASSIGN using 50 independent random number seeds were performed and converged on the assignments listed in Supplemental Table S1. Signals that were not assigned unambiguously in TDP-43-LC are listed in Supplemental Table S2. Control runs were performed to confirm that the assignments were unique. MCASSIGN run with the sequence of the His-tagged construct failed to change the assignments. MCASSIGN run with deletion of residues for which the assignments converged (365–400) from the input sequence and the same signal tables failed to converge on a unique result.

The assigned ^13^CA, ^13^CB, ^13^C, and ^15^N chemical shifts were input into the TALOS-N program using standard parameters.^31^ Residues with ‘Strong’ and ‘Generous’ torsion angle predictions were used to construct the plot in Figure 5. Error bars are the standard deviation reported by TALOS-N.

## Results

### TDP-43-LC Forms Liquid Droplets that Transition into Amyloid-Like Fibrils

As denaturant is removed from a purified solution of recombinant His-tagged TDP-43-LC through dialysis, the sample becomes turbid after approximately 0.5 h and persists for 2–3 h depending on the diameter of the dialysis tubing. The bright-field microscopy images in Figure 1a and Supplemental Video S1 reveal that the turbid solution is densely populated with spherical liquid droplets. The droplets, up to ~15 μm in diameter, are dynamic and fuse with one another when they come into contact. Figure 1b shows a fusion event for a droplet resting on the glass coverslip used in the imaging experiment. Fusion events are consistent with the liquid droplets forming through a liquid-liquid phase separation process.^8^

**Figure 1.**
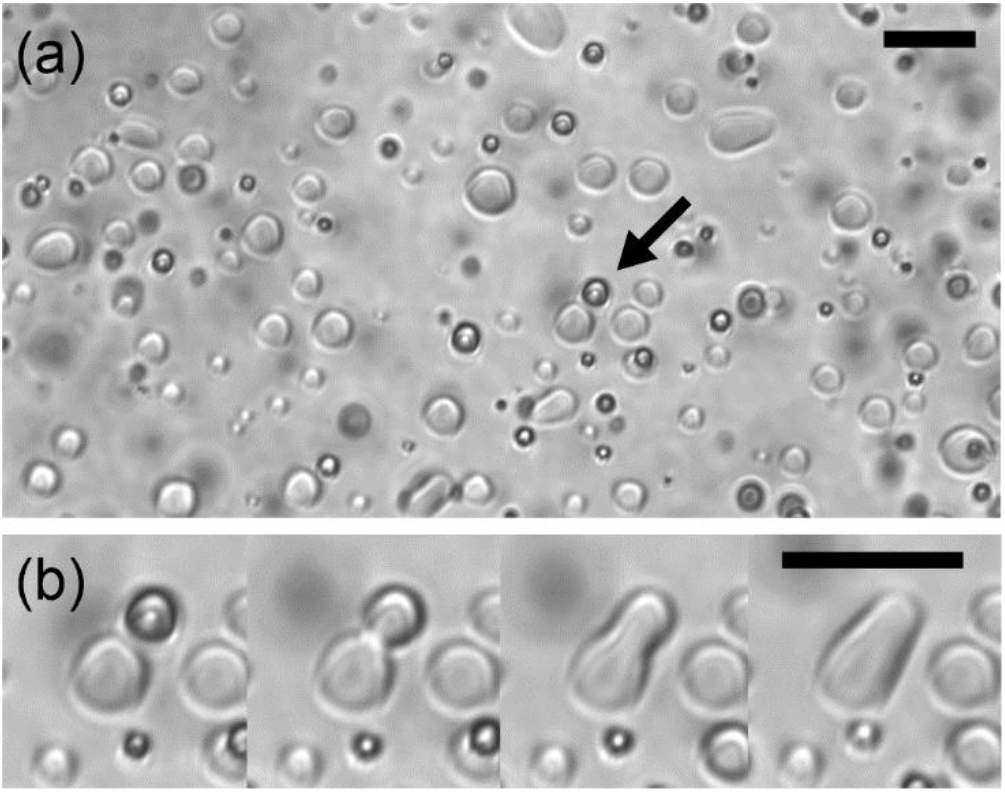
TDP-43-LC Liquid Droplets. (a) A bright field DIC microscope image of TDP-43-LC liquid droplets recorded 0.5 h after the start of dialysis. (b) Enlarged images of TDP-43-LC liquid droplets taken 1 s apart, illustrating a fusion event. The arrow in panel (a) indicates the liquid droplet undergoing the fusion event in panel (b). In both panels, the black scale bar represents a 20 μm distance.

Supplemental Figure S1 shows the quantification of the denaturant concentration under these dialysis conditions. After 1 h 75% of the urea is removed from the sample and after 3 h 97% of the denaturant is removed from the sample. Therefore, the droplets first appear when the urea concentration is approximately 1.5 M and then begin to disappear 1–2 h after the denaturant concentration is reduced to 0.2 M. This observation is consistent with other studies that show TDP-43-LC liquid droplets do not persist for extended periods of time at similar pH and electrolyte concentrations.^18,32^

Figure 2a shows that TDP-43-LC liquid droplets induce marginal increases in ThT fluorescence compared to the denatured protein. The increase is consistent with ThT entering a viscous, confined, environment.^33^ Figure 2a also shows a marked increase in ThT fluorescence as the droplets are aged for a period of 124 h, during which time the turbid solution converts to a clear solution containing a white precipitate-like material in the dialysis tubing. This larger increase in fluorescence is consistent with a ThT interaction with cross-β protein fibrils, not amorphous protein aggregation.^34^

**Figure 2.**
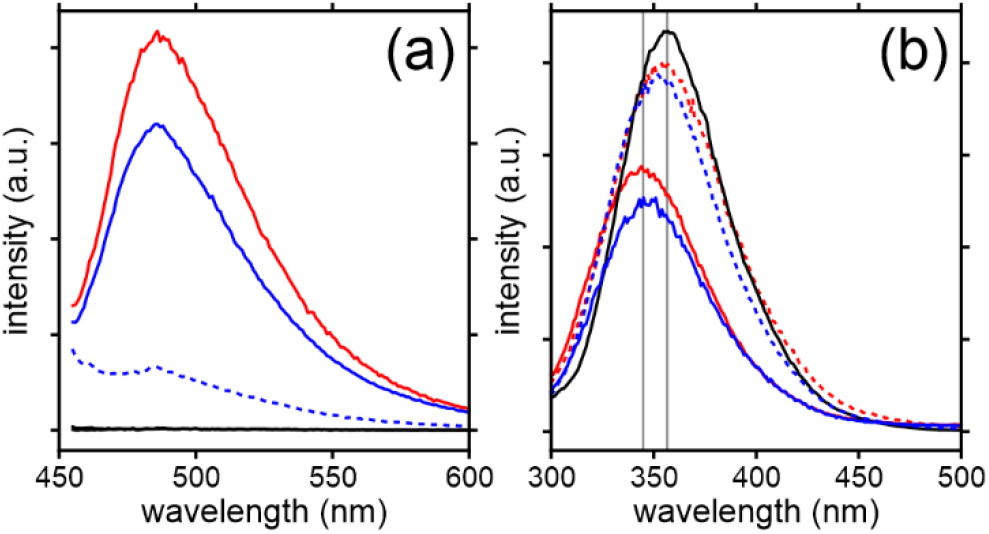
TDP-43-LC fluorescence spectra. (a) ThT fluorescence spectra of TDP-43-LC in denaturing conditions (black line), 70 min after the start of dialysis when liquid droplets are present (blue dashed line), after aging the liquid droplets for 124 h (blue solid line), and after the flash diluted sample was rotated for 24 h (red solid line). TDP-43-LC ThT fluorescence spectra immediately after flash dilution is indistinguishable from the protein in denaturing conditions (black line). (b) Trp fluorescence spectra of TDP-43-LC in denaturing conditions (black line), 70 min after the start of dialysis when liquid droplets are present (blue dashed line), after aging the liquid droplets for 124 h (blue solid line), immediately after flash dilution (red dashed line), and after the flash diluted sample was rotated for 24 h (red solid line). The vertical gray lines in panel (b) are drawn at 357 nm and 344 nm.

Figure 2b shows that the intrinsic Trp fluorescence spectrum of TDP-43-LC in liquid droplets exhibits a relatively small hypochromic shift relative to the spectrum of denatured TDP-43-LC. The fluorescence intensity peak changes from 357 nm in the denatured state to 354 nm in the liquid droplet state. The spectrum of the aged liquid droplets reveals a larger hypochromic shift with a maximum at 344 nm. These shifts are indicative of the Trp residues converting from an unfolded and solvent exposed state to a more ordered and partially buried configuration.^35^

Rapid buffer exchange, or flash dilution, from the denatured conditions for TDP-43-LC results in the immediate formation of a white precipitate-like material in the solution. Figures 2a and 2b show the ThT and Trp fluorescence spectra of this sample. The ThT fluorescence spectra of the flash diluted sample initially shows no increase in fluorescence intensity. After rotating the sample for 24 h, the fluorescence intensity increases, consistent with the formation of cross-β protein fibrils. The Trp fluorescence intensity peak of the flash diluted sample shows a minimal shift from 357 nm to 354 nm immediately after dilution of the denaturant with a further shift to 345 nm after rotation for 24 h, a net hypochromic shift, consistent with the conversion of solvent exposed Trp residues to a more ordered and less exposed conformation. These spectra show that the flash diluted sample has the same ThT and Trp fluorescence properties as the aged liquid droplets.

Figures 3c and 3d show negatively stained TEM images of the aged liquid droplet sample and the flash diluted sample, respectively. In both cases, cross-β-like fibrils with a slightly twisted appearance are observed. Significant bundling of the fibrils is observed in addition to a few dispersed single fibrils. The level of resolution present in these micrographs and the degree of fibril bundling make the fibrils indistinguishable between the two preparations.

**Figure 3.**
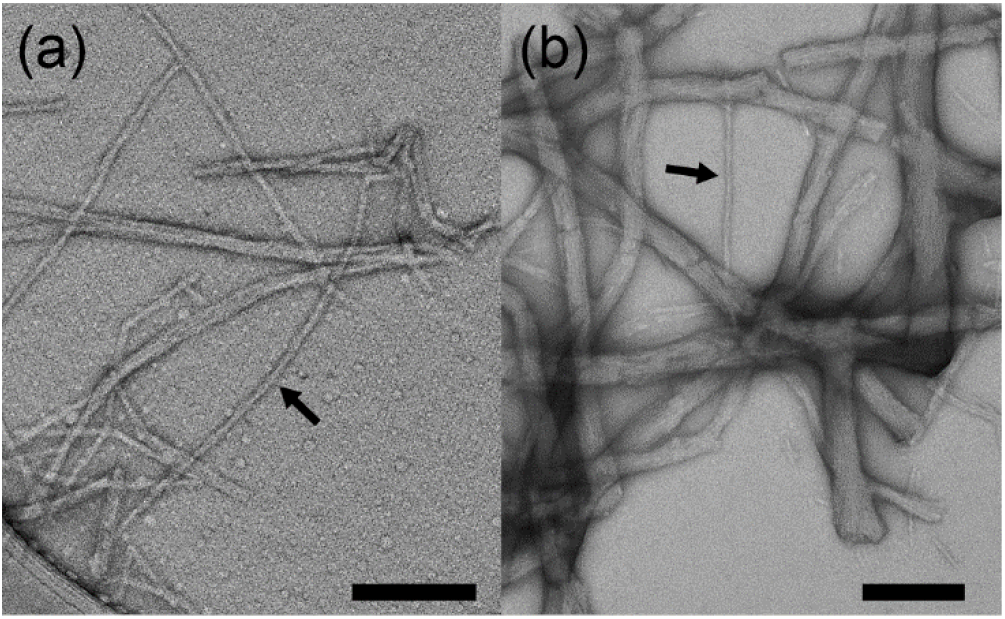
TEM Imaging of TDP-43-LC. (a) a negatively stained TEM image of TDP-43-LC aged liquid droplets. (b) a negatively stained TEM image of flash diluted TDP-43-LC. Black scale bars represent a 200 nm distance. Black arrows point to a single unbundled fibril.

### Aged Liquid Droplet and Flash Diluted TDP-43-LC Fibrils are Structurally Similar

Figure 4 shows magic angle spinning (MAS) solid state NMR spectra of the aged liquid droplet and flash diluted TDP-43-LC samples. These cross polarization (CP)^36^ based spectra, recorded at a moderate MAS frequency (13.0 kHz) with high power ^1^H decoupling (83.3 kHz), report on rigid segments of the TDP-43-LC protein in these two preparations. The aliphatic region of the 2D ^13^C-^13^C CP-DARR^37,38^ (cross polarization dipolar assisted rotational resonance) spectrum of the aged liquid droplets in Figure 4a reveals the clear presence of multiple Ser and Ala residues with ^13^C chemical shift values indicative of β-strand structure. In this spectrum, strong signals are also present from Ala residues with helical and random coil chemical shift values. There is a weak signal from one Ile residue, as well as signal intensity corresponding to the highly ambiguous region for Arg, Asp, Asn, His, Gln, Glu, Leu, Lys, Met, Phe, Trp, and Tyr residues. However, the aromatic region of the spectrum (not shown) is essentially devoid of signals.

**Figure 4.**
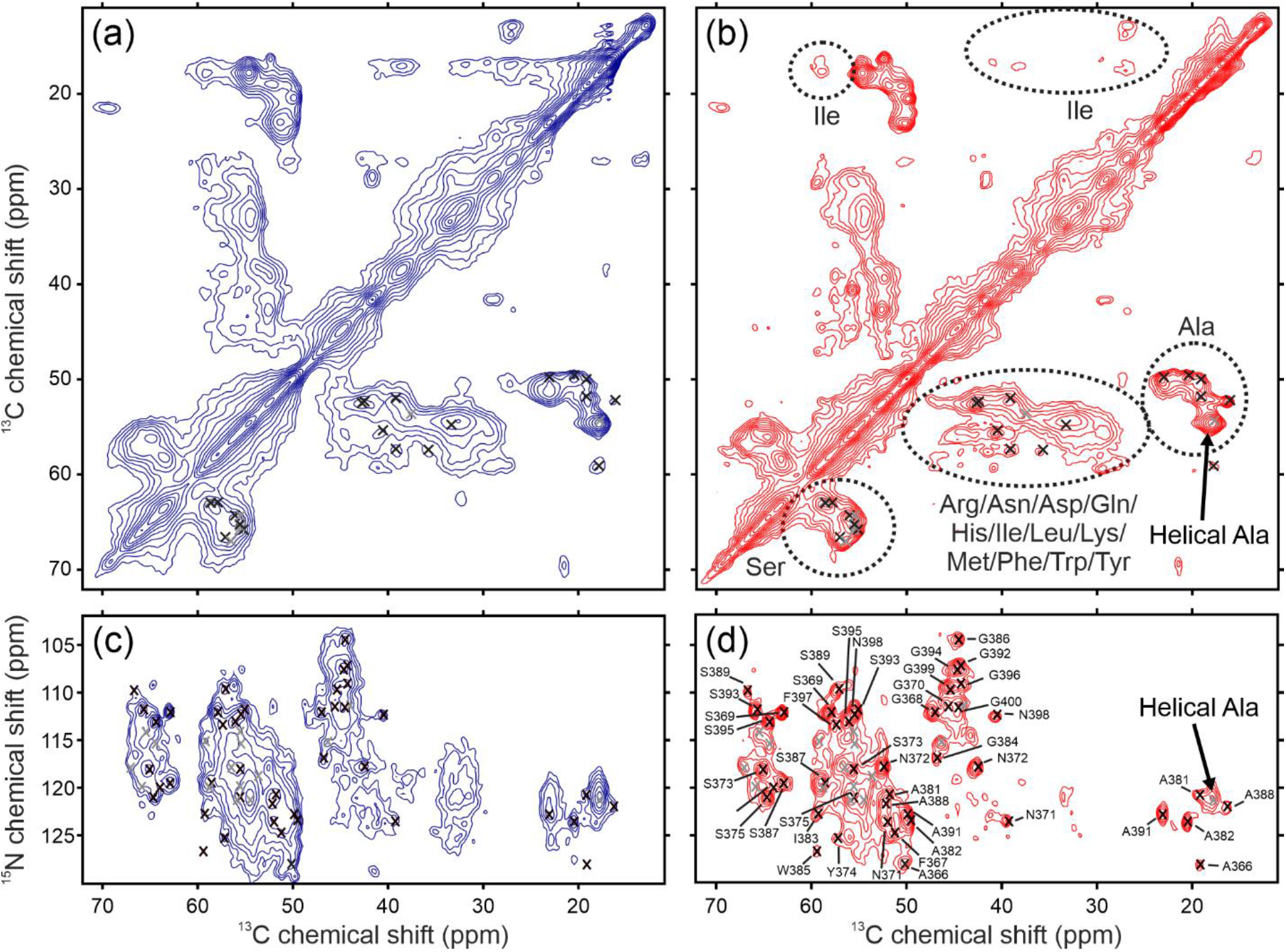
Solid State NMR spectra of TDP-43-LC. (a) aliphatic region of a ^13^C-^13^C CP-DARR spectrum of aged TDP-43-LC liquid droplets. (b) aliphatic region of a ^13^C-^13^C CP-DARR spectrum of flash diluted TDP-43-LC. (c) aliphatic region of a CP-NCACX spectrum of aged TDP-43-LC liquid droplets. (d) aliphatic region of a CP-NCACX spectrum of flash diluted TDP-43-LC. The residue-type assignments shown in panel (a) are based on statistical values of protein chemical shifts. X-marks indicate the locations of signals observed in the 3D spectra presented in this paper, colored black for unambiguously assigned signals and colored gray for unassigned signals. In panel (d), the residue specific assignments are those determined in this paper. Successive contours are drawn in intensity increments of 1.4 for panels (a) and (b) and increments of 1.3 for panels (c) and (d).

Figure 4b shows the aliphatic region of the 2D ^13^C-^13^C CP-DARR spectrum of the flash diluted TDP-43-LC protein, recorded under the same conditions as the aged liquid droplet sample. The spectrum contains resonances and resonance intensities indicating that the same amino acid types are immobilized in the fibrils formed through the flash dilution method as are in fibrils formed by aging of liquid droplets. In addition, the positions of the resolved resonances and resonance envelopes are highly similar. The data indicate that the underlying structure of the fibrils formed by the two sample preparation methods are nearly identical. Supplemental Figure S2 shows an overlay of the ^13^C-^13^C CP-DARR spectrum of these two samples, further illustrating the high degree of similarity between them.

Figures 4c and 4d show the aliphatic region of 2D CP-based NCACX spectra of the aged liquid droplet and flash diluted TDP-43-LC, respectively. These spectra are also recorded using a moderate MAS frequency (13.0 kHz) and high power ^1^H decoupling (83.3 kHz), therefore reporting on residues immobilized in the fibril core. The spectra exhibit well-resolved signals with ^13^C and ^15^N linewidths of ~1.0 ppm, and have well dispersed ^15^N chemical shift frequencies, consistent with a well-ordered fibril core. The presence of numerous Gly residues is also evident from these spectra. Supplemental Figure S3 shows an overlay of these spectra, reinforcing their similarities. Supplemental Figure S4 shows 2D CP-based NCOCX spectra of the aged liquid droplet and flash diluted TDP-43-LC. These spectra have the same pattern of NMR signal intensity, also indicating that the fibrils formed through the two sample preparation methods are constructed from TDP43-LC proteins in highly similar conformations.

The spectra of the aged liquid droplets appear to have similar linewidths as the flash diluted fibrils, although a broad intensity underlies the sharper signals. Global ^1^H T_1ρ_ measurements obtained with a ^1^H-^13^C CP pulse sequence and ^15^N T_2_ measurements obtained with a CP-NCA pulse sequence reveal that the relaxation parameters for non-glycyl CA sites are not significantly different between the two sample preparations. Glycine CA signals were integrated separately in the ^15^N T_2_ analysis and gave similar values, although there was overlap with CB signals of other residues (based on the assignments determined in this paper). For the aged liquid droplets and the flash diluted fibrils, the ^1^H T_1ρ_ measurements revealed biexponential decays with apparent decay constants of 1.0/6.6 ms and 1.7/8.3 ms respectively, and the ^15^N T_2_ measurements revealed monoexponential decays with decay constants of 9.0 ms and 8.4 ms respectively. These relaxation measurements are consistent with a highly ordered fibril core for both samples and are also consistent with the ^15^N linewidths observed for resolved resonances.

Supplemental Figures S2 and S3 show the ^13^C-^13^C CP-DARR and NCACX difference spectra for the aged liquid droplet and flash diluted TDP-43-LC. Residual signals in the difference spectra arise from immobilized sites in the aged liquid droplet spectrum that are not present in the flash diluted fibril spectrum. Signal intensity from Ala, Gly, and Ser residues are present in these spectra. The residual Gly and Ser signals have different chemical shifts than those from the fibril core region, indicating these residues likely have different conformations and chemical environments than those composing the fibril core. The Ala regions of the difference spectra have some intensity with the same chemical shifts as the flash diluted spectra for helical, β-strand, and random coil Ala residues. This intensity indicates either that some of the Ala residues are even more strongly ordered in the aged liquid droplets than in the flash diluted sample or that additional Ala sites with nearly identical chemical shifts as those in the flash diluted sample have become ordered in the aged liquid droplet sample. However, three of the six easily distinguished Ala signals are absent from the difference spectra. Except for some additional broad signal intensity from Gln residues, the difference spectra are devoid of signals from any other amino acids, such as the ten Met residues in TDP-43-LC.

Together the relaxation measurements and the difference spectra show that the underlying intensity in the aged liquid droplet sample is consistent with reduced molecular motions for portions of TDP-43-LC in heterogeneous conformations, not increased dynamics of the rigid fibril core in the sample. The extra intensity in the spectra of the aged liquid droplets is therefore either due to dynamic unstructured regions outside the fibril core becoming immobilized as the liquid droplet age, or incomplete conversion of all TDP-43-LC molecules in the sample into the same fibril conformation. The distribution of Gly, Ala, and Ser residues throughout the entire TDP-43-LC sequence prevents sequence-specific assignment of this extra intensity. Despite the subtle differences in the spectra of the aged liquid droplet sample and the flash diluted sample, the identical chemical shifts for the well-ordered sites show that both sample preparation methods yield fibrils constructed from TDP-43-LC in similar molecular conformations. In further support of this interpretation, the TEM images in Figure 3 do not show any obvious differences in fibril morphology between the two samples.

### Structural Characterization of the Fibril Core

3D CP-based NCACX, NCOCX, and CANCO spectra were recorded on the flash diluted sample using similar experimental conditions as the 2D spectra already presented. These spectra exhibited large signal-to-noise ratios and allowed resolution of 40, 39, and 43 unique signals, respectively. Supplemental Figures S5, S6, and S7 show 2D planes from these spectra, illustrating the resolution and signal-to-noise of the 3D spectra. A computationally assisted assignment procedure^30^ resulted in statistically significant and unambiguous sequence-specific chemical shift assignments for residues Q365–G376 and G380–G400. These assignments indicate that the rigid core of the TDP-43-LC fibrils is composed of one Gln, five Ala, two Phe, eleven Gly, seven Ser, four Asn, one Tyr, one Ile, and one Trp. The computational algorithm ensures that the sequence specific assignments are unbiased and unambiguously unique.^39^ A robust computational method is essential due to the strong bias of the TDP-43-LC sequence toward Gly, Ser, Asn, and Ala residues, which are 25%, 16%, 14%, and 9% of the total amino acid content, respectively. The observation that 82% of the NMR signals are one of these four amino acid types further reinforces the need for computationally assisted assignments. Very strong evidence for the uniqueness of the assignments is provided by the convergence of the algorithm on an identical result for 50 independent calculations with 50 independent random number seeds.

Signals for three Ser, one Ala, one Gly, and one ambiguous-type residue were not assigned. Two Ser residues and the ambiguous-type residue can be tentatively but not unambiguously assigned to residues S377–S379. The lone unassigned Ala signal is quite strong in all 3D spectra, suggesting it may derive from more than one Ala residue in the protein. Signals in the NCOCX data suggest the residue giving rise to this signal is part of an Ala-Ala pair in the protein sequence and may derive from more than one site in the protein. The only remaining Ala-Ala pairs in the unassigned region of TDP-43-LC are the five Ala residues within residues A324–A329. The chemical shift values for this signal are 54.7 ppm and 17.9 ppm for CA and CB, respectively, indicating the residue (or residues) giving rise to this signal intensity is in a helical conformation. The other remaining unassigned signals are not present in all three of the 3D spectra and are of relatively low signal-to-noise. The computational assignment procedure failed to assign any residues to the His-tag region of the TDP-43-LC construct, indicating this region is not structured in the aged liquid droplet or flash diluted fibril samples. Assignment calculations run using an input sequence with the residues after Q365 deleted failed to converge on unique or unambiguous assignments for any residue, indicating that the observed signals are not consistent with the amino acid sequence prior to residue Q365. Disorder in the N-terminal region is further supported by the lack of any strong His, Pro, Met, and Thr signals in the CP-based NCACX and NCOCX spectra.

Figure 5a shows ^1^H-^13^C refocused INEPT spectra of the flash diluted fibrils and the aged liquid droplets. The flash diluted sample exhibits relatively weak signals and amino acid type assignments are difficult for this spectrum. However, the presence of aromatic signals consistent with His sidechain sites suggests the N-terminal region containing the His-tag in our TDP-43-LC construct is dynamic and disordered. The relatively weak signals also suggest that only a small portion of the LC domain is highly mobile in the sample, as other LC domain fibrils with substantial regions outside the immobilized core exhibiting rapid dynamics and significant disorder give rise to much stronger signals in these experiments.^11^ The ^1^H-^13^C INEPT spectrum of the aged liquid droplet sample only contains strong signals that derive from highly dynamic methyl groups. Together with the CP based data collected on these samples, the INEPT spectra suggest that the regions outside the highly immobilized fibril core segment sample heterogenous conformations with limited molecular motion but have not adopted a singular conformation.

**Figure 5.**
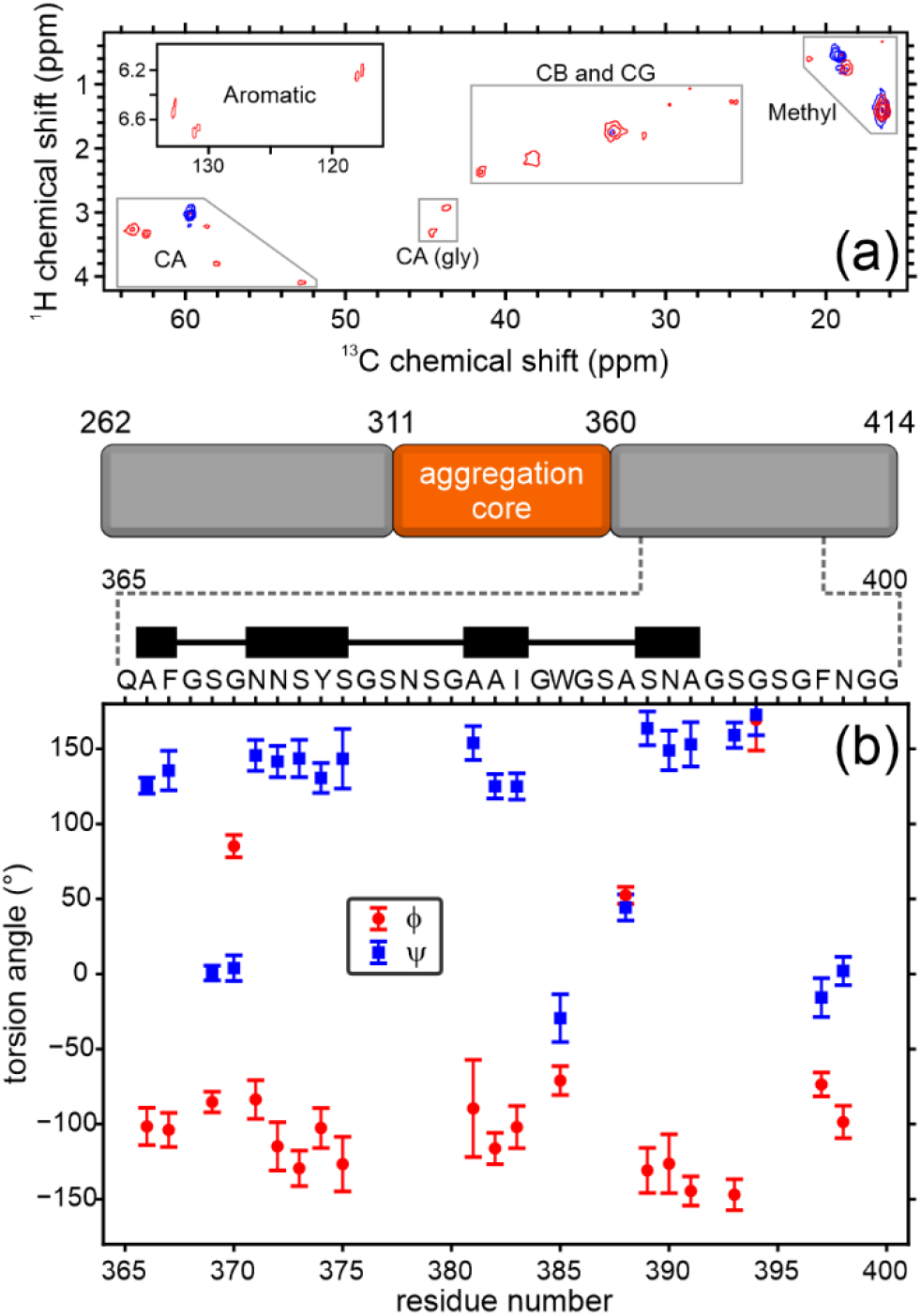
Structure and Domain Organization in TDP-43-LC. (a) ^1^H-^13^C INEPT spectrum of TDP-43-LC. The spectrum for the flash diluted sample is drawn in red, and the spectrum for the aged liquid droplet sample is drawn in blue. Contours are drawn with a factor of 2.0 between levels. (b) TALOS-N torsion angle prediction as a function of residue number. The ‘Strong’ and ‘Generous’ sites are plotted, and the error bars represent the standard deviation output by the TALOS-N program. The letters at the top of the plot indicate amino acid types. The thick lines in the black block diagram represents the regions predicated to have β-strand conformations. The cartoon depicts the location of the residue 311–360 aggregation core in TDP-43-LC. The gray dashed lines highlight the core region characterized in this study.

Figure 5b shows the TALOS-N torsion angle predictions based on the observed and assigned ^13^C and ^15^N chemical shifts. Residues A366–F367, N371–S375, A381–I383, S389– A391, and S393 are predicted to have φ/ψ torsion angles indicative of extended conformations consistent with β-strand secondary structure. Residues S369–G370, W385, A388, G394, and F397–N398 have φ/ψ torsion angles indicative of turn-like conformations. Signals were observed for G368, G376, G380, G384, G386–S387, G392, S395–G396, and G399–F401 but TALOS-N failed to predict φ/ψ torsion angles for these sites. These predictions indicate there are at least three segments in β-strand conformations in the TDP-43-LC fibrils.

## Discussion

### TDP-43-LC Liquid Droplets Mature into β-strand-Rich Protein Fibrils

The primary result from the measurements presented in this paper is the identification of the segment of TDP-43-LC that condenses into a stable well-defined structure upon the aging of liquid droplets. Specifically, we have found that (i) TDP-43-LC converts into a ThT positive fibril structure upon the removal of denaturant (Figure 2 and 3), (ii) the liquid droplet state is a distinct intermediate between the denatured and fibril state (Figure 1 and Figure 2), (iii) residues 365 to 400 form the rigid core of the fibrils (Figures 4 and 5), (iv) the regions of TDP-43-LC flanking the fibril core are not highly dynamic (Figure 5), and (v) the fibril core is constructed from at least three segments of the TDP-43-LC in β-strand conformations with a dynamic loop between residues 376 and 380 (Figure 5).

Our results clearly show that liquid droplets of TDP-43-LC mature into β-strand rich fibrils stabilized by residues 365–400. Support for our conclusion comes from a study by Babinchak *et al.* on a similar TDP-43-LC construct (His-tagged residues 274–414).^32^ First, atomic force microscopy images clearly show fibrils emanating from dried liquid droplets.^32^ Second, although this study used reduced pH to slow the liquid droplet to fibril transition, fibrils with an identical appearance and significant clumping were observed at near neutral pH.^32^ Third, TDP-43-LC liquid droplets initially exhibit minimal ThT fluorescence followed by a significant increase in intensity after a lag phase.^32^ Finally, fibrils obtained without the formation of liquid droplets had the same visual appearance and fluorescent properties as those nucleated in the liquid droplets.^32^ The similarities between our sample preparations and those in this study strongly support our interpretation that the fibrils characterized here are the structure that condenses out of the liquid droplets.

### Competing Structural Conformations in TDP-43-LC

In the context of existing studies on the C-terminal LC domain of TDP-43, our results are quite intriguing. Residues 311–360 of TDP-43-LC are necessary for aggregation in cells^17^ and in isolation, this ‘aggregation core’ segment forms stable cross-β protein fibrils through backbone β-strand hydrogen bonding, sidechain Gln polar zippers, and a substantial hydrophobic core^20,21^ reminiscent of those formed by the β-amyloid peptide^40^ and α-synuclein.^41^ Notably, the cryo-EM structures of the residue 311–360 fragment indicate extensive hydrophobic interfaces involving the sidechains of five Met residues and five Gln residues positioned to facilitate intermolecular hydrogen bonds.^21^ Yet after aging liquid droplets formed by the complete LC domain, we find the most stable segment of the protein begins at least ten residues after this region. This ‘second core’ region lacks any structured Met residues, has a single weakly structured Gln site, contains eight more Gly and four more Ser residues. These differences indicate the ‘second core’ has limited interactions between extended sidechain moieties and a substantially smaller fraction of buried hydrophobic surface area. These differences seem to suggest the second core might be less thermodynamically favored than the aggregation promoting core between residues 311–360. However, our data indicate the ‘second core’ is the stable conformation for aged liquid droplets of TDP-43-LC.

Solution NMR measurements indicate that residues 321–341 transiently populate a helical conformation in TDP-43-LC monomers.^18^ Interactions between the helical region and residues 382–385 (squarely in our ‘second core’) facilitate the formation of oligomeric species in a concentration dependent manner.^18^ The liquid droplet state is difficult to probe using solution NMR due to rapid rigidification of TDP-43-LC droplet samples.^18^ However, mutations that disrupt the helical structure correlate with a decreased propensity to form liquid droplets.^18^ Both transient helix formation by residues 321–341 and their brief interactions with residues 382–385 seem to be essential for liquid droplet formation.^18^ It should not escape notice that the unassigned Ala residue (or residues) in our data has a chemical shift signature indicating it is in a helical conformation, although the chemical shift values are not precisely those observed by solution NMR^18^ for the multiple Ala residues in the helical prone region. We were not able to unambiguously assign this residue due to a lack of strong magnetization transfer to neighboring residues. However, the data from our measurements are consistent with part of the poly-Ala stretch between residues 324–329 adopting a helical-like conformation that is loosely ordered and only immobilized sufficiently to permit CP based magnetization transfers for one or two Ala sites in this region and no other residues.

Our data are therefore consistent with an assembly process where the helical region between residues 321–341 is prevented from converting into its fibril conformation through a weak interaction with the primarily β-strand structure formed by residues 365–400. We propose that our solid state NMR data from fibrils of TDP-43-LC, the cryo-EM^21^ and solid state NMR^20^ characterization of fibrils formed by residues 311–360, and the solution NMR and mutational analysis^18^ of TDP-43-LC are all consistent with a model where a functional propensity for residues 365–400 to form stable β-strand structures prevents detrimental aggregation of residues 311–360.

A recent solid state NMR study on TDP-43-LC aggregates seeded from protein isolated from *E. coli* inclusion bodies shows that both core regions can form a stable fibril conformation simultaneously (i.e. residues 306–343 and 362–397).^42^ However, our result is quite distinct from that work. First, our samples were purified under conditions that completely solubilized the inclusion body fraction of the *E. coli*. Second, the chemical shifts we report are significantly different for 24 out of the 29 structured residues between 365–397, as defined by a greater than 0.5 ppm difference in CA or CB chemical shifts. These differences, illustrated in Supplemental Figure S8, are not explained by a constant additive correction to all shifts or uncertainties due to low signal to noise. The fibril structure stabilized by the *E. coli* inclusion body environment is therefore different than what matures out of TDP-43-LC liquid droplets in our hands. One likely explanation for this difference is the lack of a templating structure stabilized by the molecular components of the *E. coli* inclusion body environment. This longer ordered segment incorporating both fibril-prone core regions could be the TDP-43-LC structure after the helix in residues 321–341 fully converts to its β-strand conformation. However, this interpretation requires that the fibril structures adopted by both the ‘second core’ segment identified here and the aggregation core in residues 311–360 undergo significant structural changes, since the boundaries of the rigid segments are not the same and the chemical shifts for the ordered residues in common between these preparations are different. Supplemental Figure S8 shows the chemical shift differences for all residues that are structured in either the aggregation core (residues 311–360) or the ‘second core’ (residues 365–400), and the *E. coli* inclusion body fibrils.

### TDP-43-LC Domain Interactions in the Full-Length Protein

Solid state NMR measurements from Shenoy *et al*.^19^ on fibrils formed by the full-length TDP-43 sequence, residues 1–414, and a set of truncation mutants provide context for the LC domain assembly process when the N-terminal RRM and oligomerization motifs are present. The presence of multiple Ala residues in both fibril-prone regions provides a straightforward way to compare these spectra^19^ with our data and the data from the residue 311–360 fibrils^20^. Figure 6 outlines these differences. Figure 6c shows that fibrils formed by the full-length TDP-43 sequence do not give rise to strong Ala signals with chemical shifts characteristic of β-strand conformations.^19^ A truncated version of TDP-43-LC missing residues 367–414 gives a similar pattern of Ala signals as the full-length protein.^19^ However, the helical Ala signal present in our spectra is also observed in these spectra. Figure 6c also shows that a similar construct as the one investigated in this paper, His-tagged residues 267–414, gives rise to a pattern of Ala signals^19^ partially consistent with our spectra. Signals for the helical Ala, A381, and A382 are present, but there are three peaks in the vicinity of A391 and signals for A366 and A388 are absent. The differences are likely due to the different electrostatic environments and protonation states of these samples (in reference ^19^ the fibrils were prepared in 20 mM MES buffer at pH 7.5 without salt and extensively washed with water, while the fibrils in our study are prepared in 20 mM sodium phosphate and 200 mM sodium chloride at pH 7.4). Figure 6b shows the chemical shifts from the residue 311–360 fibrils.^20^ Comparison with our spectra (Figure 6a) and the that of the full-length TDP-43 (Figure 6c)^19^ reveal that this conformation is not strongly ordered in the fibrils of longer TDP-43 constructs. Figure 6d shows the chemical shifts from the *E. coli* inclusion body templated TDP-43-LC fibrils. Comparison with the Ala signals from other TDP-43 fibrils (Figures 6a-c) further illustrates the structural differences for these templated fibrils.

**Figure 6.**
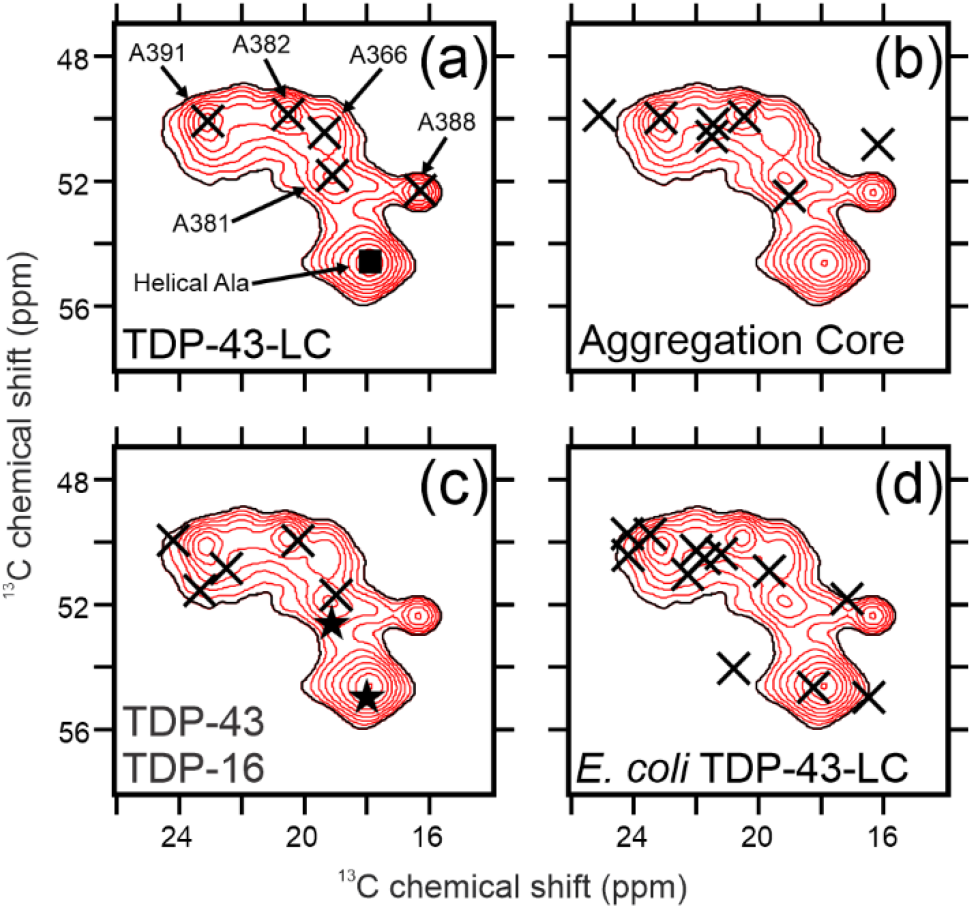
Comparison of TDP-43 Solid State NMR Results. Each panel shows the Ala signal intensities from the residue 262–414 flash diluted fibrils from this paper in red. In panel (a) the X-marks indicate the assigned Ala signals and the black square indicates the unassigned helical Ala signal from the data presented in this paper. In panel (b) the X-marks indicate the assigned Ala signals from the fibrils formed by the aggregation core, residues 311–360, from reference ^20^. In panel (c) the X-marks and stars indicate the locations of Ala signal intensities from the fibrils formed by TDP-16, residues 274–414, from reference ^19^. The stars indicate the Ala peaks that are present in the spectra of the full-length TDP-43 fibrils from reference ^19^. In panel (d) the X-marks indicate the assigned Ala signals from the *E. coli* inclusion body seeded residue 274–414 fibrils from reference ^42^.

What does it mean that the resonances reporting on the fibril conformations for residues 311–360 and 365–400 are absent from the spectrum of the full-length TDP-43 fibrils? It is likely that in the full-length TDP-43 CP based spectra increased molecular motion of the protein backbone in the fibril-prone region leads to poor magnetization transfers and reduced or missing signal intensities. For instance, the β-strand-rich segment we identified as 365–400 may adopt a dynamic β-strand conformation in full-length TDP-43 fibrils that is mobile enough to not give rise to signals. Despite this difference, the full-length fibrils^19^ and our samples both exhibit the intense Ala signal we report as helical structure in residues 324–329. The presence of this same strong helical Ala signal suggests that both structures lack even a loosely ordered β-strand-rich conformation from residues 311-360. However, solid state NMR and cryo-EM studies on the residue 311–360 TDP-43-LC fragment show these residues adopt a rigid β-strand-rich structure.^20,21^ This observation reinforces our assertion that the propensity of residues 365–400 to adopt a fibril conformation directly prevents the pathological aggregation of the remainder of the TDP-43-LC. In this model, removal of residues after 367 favors the pathological assembly process of residues 311-360. This type of dynamically structured LC domain is not unprecedented, as we have recently observed such a phenomena in a LC domain of intermediate filament protein assemblies.^43^ These data all reinforce our interpretation that delicately balanced interactions determine the functional and pathological assembly of TDP-43-LC.

### LC Domains and (a Lack of) Fibril Polymorphism

Given the reduced amino acid complexity of the TDP-43-LC, it is perhaps not surprising that several segments are able to form stable fibrils. 63% of the LC domain sequence is composed of Ala, Asn, Gly, and Ser residues. Based on similarities with prion-like domains, regions rich in these residues are expected to adopt fibril conformations.^44^ Pathogenically associated amyloid-like protein fibrils such as β-amyloid and α-synuclein are highly prone to structural polymorphism.^22^ TDP-43 is found in pathological inclusions in patient tissues,^15^ and residues 285-331, in addition to residues 311-360 and 365-400, have also been observed to form stable fibrils^21^, supporting a purely pathogenic role for the TDP-43-LC. On the other hand, the full length LC domains studied to date are notably not polymorphic with respect to fibril formation.^11,23,45^ However, the lack of polymorphism observed in the fibrils formed by complete LC domains does not exclude the presence of multiple fibrillation-prone regions in these sequences. For example, electron diffraction studies on microcrystals of small fragments from outside the core fibril forming region of the FUS LC show these fragments adopt similar conformations as those that stabilize the fibrils of the complete LC domain.^7,24^ Experiments on longer segments of LC domains also support the presence of multiple fibril cores and suggest that the entire LC domain is important for understanding the assembly of the full-length proteins that contain them. For example, deletion of the N-terminal 111 residues of the FUS protein stabilizes fibrils formed by the C-terminal half of the LC domain.^25^ Our identification of a ‘second core’ outside of the aggregation core region in the complete TDP-43-LC bolsters evidence for the presence of delicately balanced competing fibril-prone regions as a common theme in LC domains.

## Conclusion

Pathological and functional assembly of the TDP-43 protein continues to be an intriguing process. Our contribution shows an interplay between competing molecular associations in TDP-43-LC allow for its phase separation into liquid droplets and the eventual condensation into β-strand rich protein fibrils. We suggest a model where loose interactions between helix-prone regions and β-strand-prone regions facilitate liquid droplet assembly with the fibril-prone region between residues 365-400 ultimately being the most favored conformation adopted by the complete LC domain. Conversion the LC domain into the pathological fibril conformation, likely involving residues 311–360, must require the catalyst of a template structure or other external forces. Given that almost all ALS and FTD related disease mutations cluster in the core fibril-prone regions^6^, it is likely these alterations rebalance the stabilizing interactions in the fibril-prone regions of TDP-43-LC. Further investigation is required to elucidate the effect of these mutations on the TDP-43-LC assembly process and to confirm if this model is representative of what occurs in living cells.

## Acknowledgements

We would like to thank the National Science Foundation and the National Institutes of Health for support to the UC Davis NMR Core Facility though grants NSF DBI-0722538 and NIH 1S10RR013871-01A1. The University of California, Davis, provided generous start-up funding for the Murray Laboratory. We also extend our sincerest and most humble thanks to Dr. Steven L. McKnight for providing the His-tagged TDP-43-LC plasmid used in this work.

