## Supplemental Figures and Captions for "Identification of the Rigid Core for Aged Liquid Droplets of the TDP-43 Low Complexity Domain"

**Supplemental Video 1.** TDP-43-LC liquid droplets after 30 min of dialysis. Each frame is taken 1 s apart and the video covers 50 s real time.

**Supplemental Figure 1. Quantification of denaturant removal during dialysis.** The black squares are derived from integrating the $^{1}H$ urea peaks from a 1D solution NMR spectrum. Details of the measurements are presented in the Experimental Section of the main text. The figure inset and the black dashed line show the best-fit exponential decay curve with a fixed initial amplitude of 6.00.

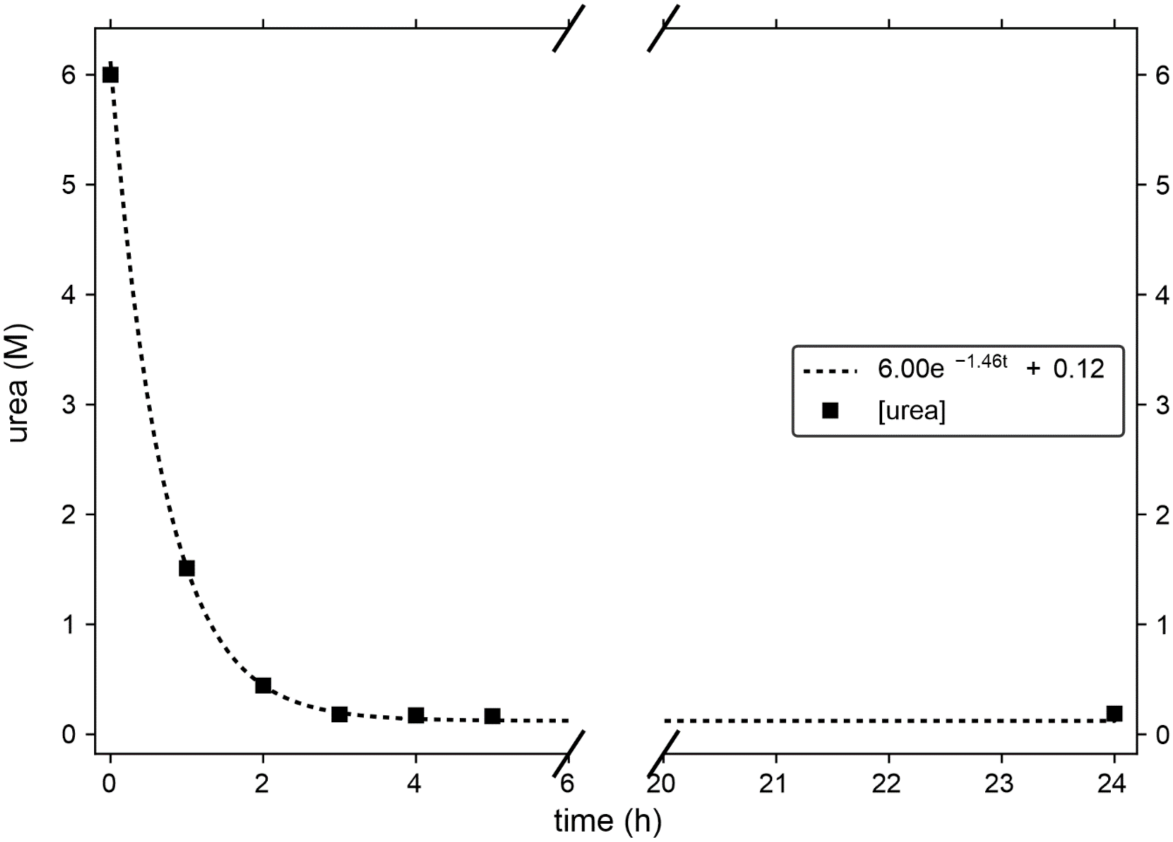

**Supplemental Figure 2. Comparison of aged liquid droplet and flash diluted TDP-43-LC $^{13}C$-$^{13}C$ CP-DARR spectra.** The left panel shows the overlay of the aliphatic regions of a $^{13}C$-$^{13}C$ CP-DARR spectra of aged TDP-43-LC liquid droplets (blue) and flash diluted TDP-43-LC (red). Black X-marks indicate the positions of signals observed in the 3D NCACX spectrum of the flash diluted TDP-43-LC. The right panel shows the difference spectrum with red X-marks indicating the positions of the X-marks from the spectrum on the left. The 1D spectra shown with the spectrum on the right are extracted from the locations of the dotted lines. Successive contour levels are drawn in intensity increments of 1.4.

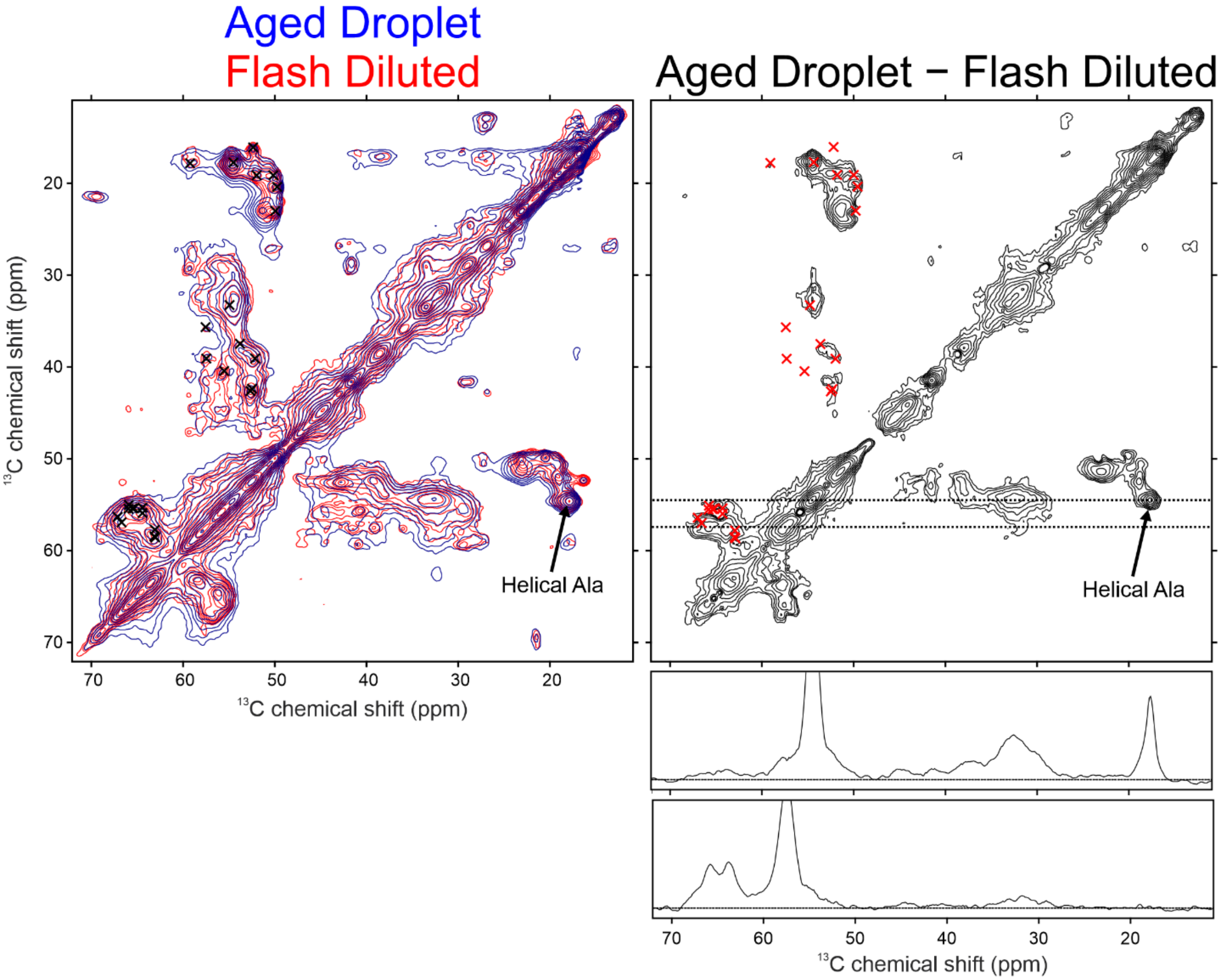

**Supplemental Figure 3. Comparison of aged liquid droplet and flash diluted TDP-43-LC CP-NCACX spectra.** The top panel shows the overlay of the aliphatic regions of the CP-NCACX spectra of aged TDP-43-LC liquid droplets (blue) and flash diluted TDP-43-LC, (red). The bottom panels show the difference spectra with red X-marks indicating the positions of peaks in the flash diluted NCACX spectrum. The 1D slices shown with the bottom spectra are extracted from the locations of the dotted lines in the 2D difference spectrum and illustrate negligible intensity for residue S373. Successive contour levels are drawn in intensity increments of 1.3.

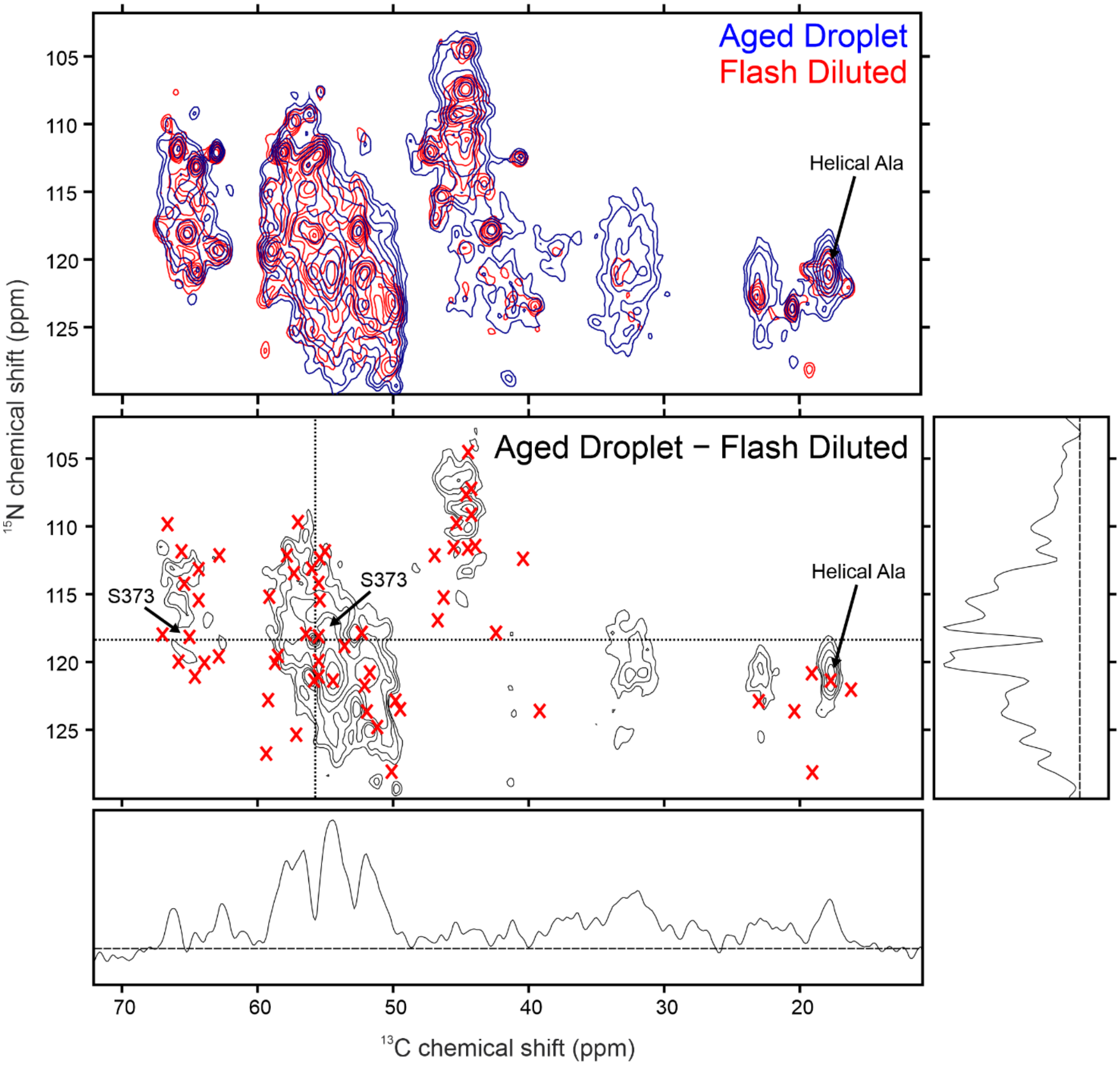

**Supplemental Figure 4. Aged liquid droplet and flash diluted TDP-43-LC CP-NCOCX spectra.** The top panel shows the aliphatic region of a 2D CP-NCOCX spectra of aged TDP-43-LC liquid droplets (blue) and the bottom panel shows the same region of the 2D CP-NCOCX spectrum from the flash diluted TDP-43-LC (red). Successive contour levels are drawn in intensity increments of 1.3. Black X-marks indicate the locations of peaks sequence specifically assigned in this paper. Gray X-marks indicate the locations of peaks not assigned in this paper.

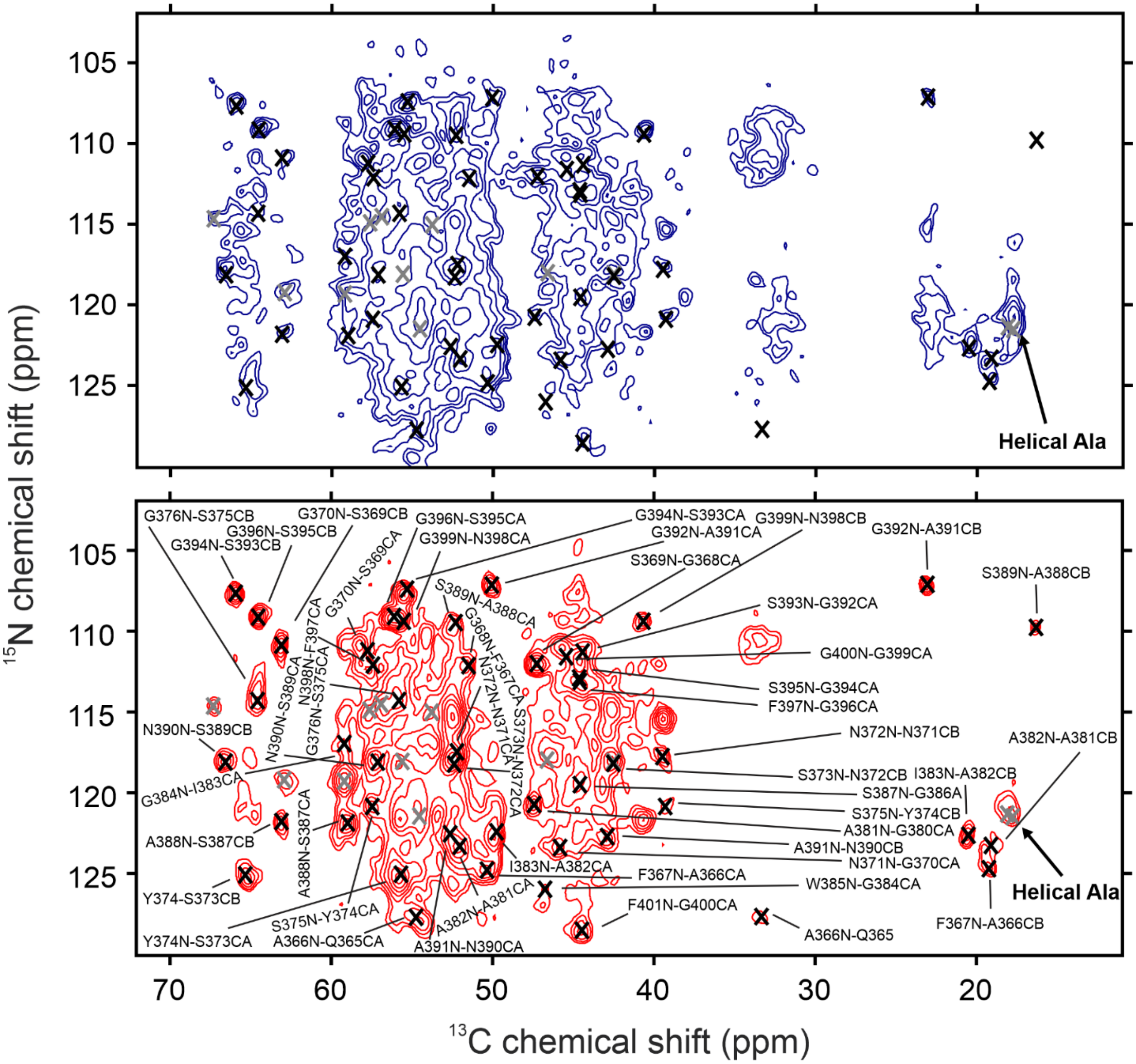

**Supplemental Figure 5**. **3D CP-NCACX spectra from flash diluted TPD-43-LC.** 2D planes from the 3D CP-NCACX spectra of flash diluted TDP-43-LC. Successive contour levels are drawn in intensity increments of 1.3. Large font labels indicate peaks that are centered in the plane and smaller font labels indicate peaks centered on neighboring planes. Grey X-marks indicate signals that are not sequence specifically assigned.

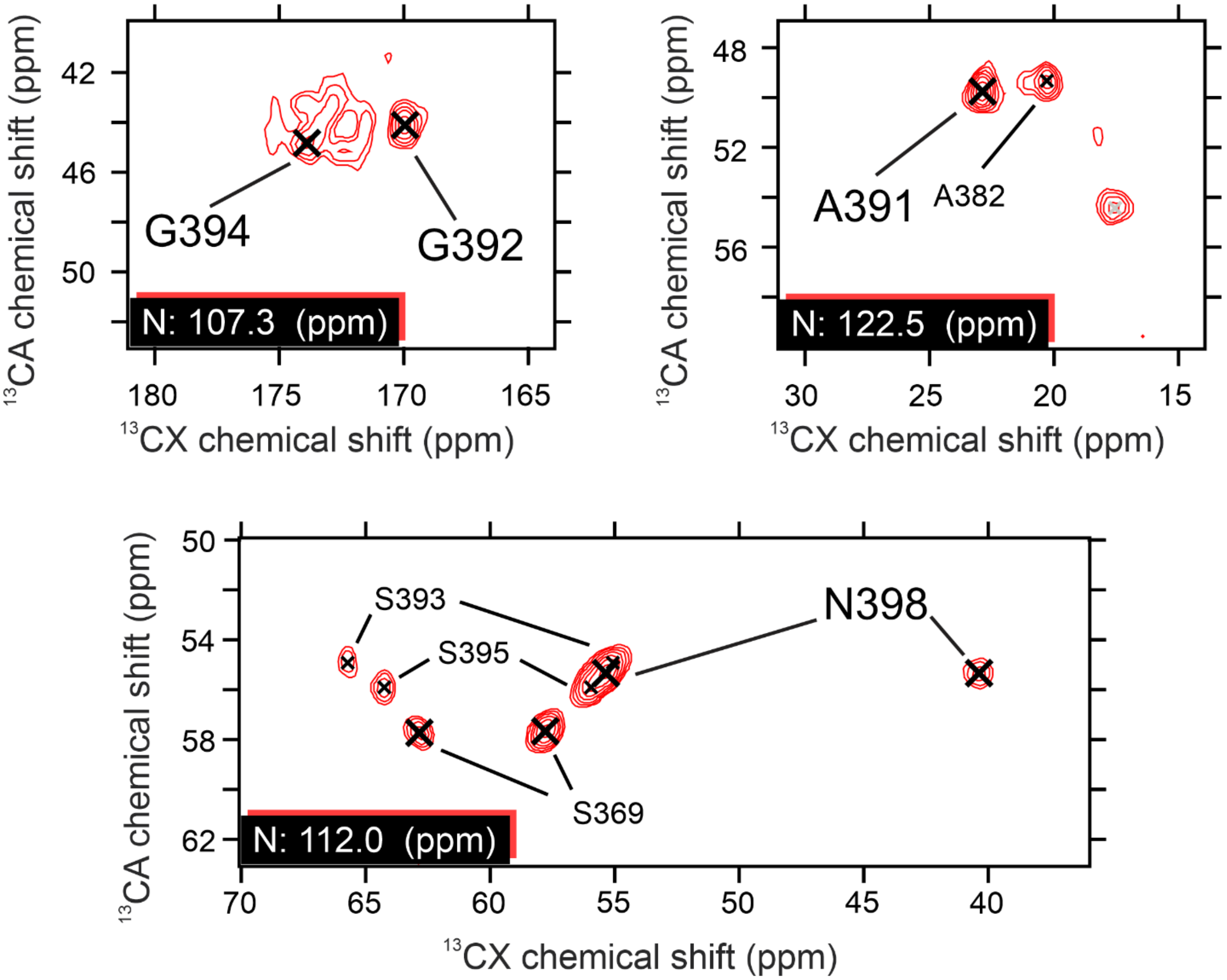

**Supplemental Figure 6**. **3D CP-NCOCX spectra from flash diluted TPD-43-LC.** 2D planes from the 3D CP-NCOCX spectra of flash diluted TDP-43-LC. Successive contour levels are drawn in intensity increments of 1.3. Large font labels indicate peaks that are centered in the plane and smaller font labels indicate peaks centered on neighboring planes. Grey X-marks indicate signals that are not sequence specifically assigned.

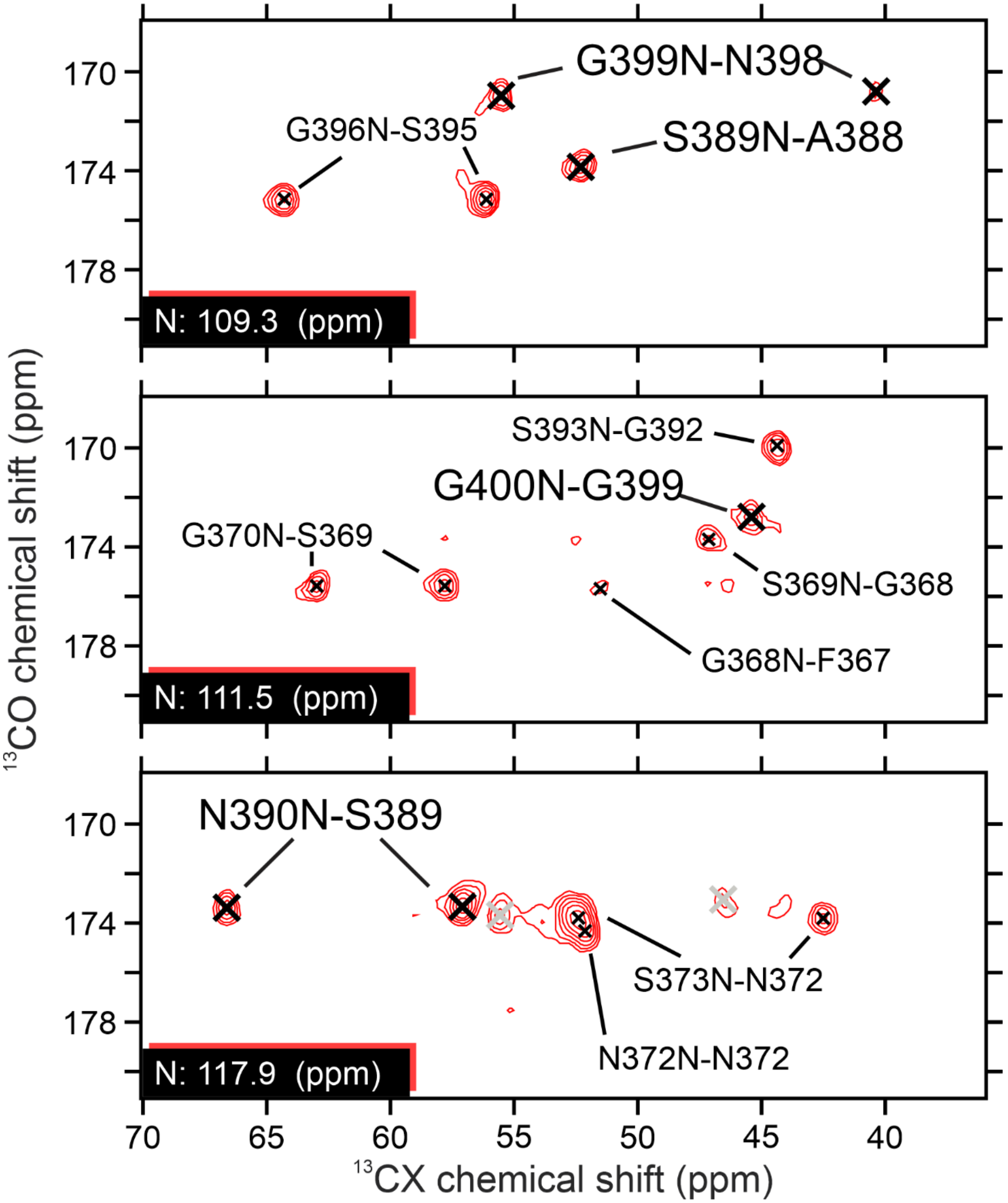

**Supplemental Figure 7**. **3D CP-CANCO spectra from flash diluted TPD-43-LC.** 2D planes from the 3D CP-CANCO spectra of flash diluted TDP-43-LC. Successive contour levels are drawn in intensity increments of 1.3. Large font labels indicate peaks that are centered in the plane and smaller font labels indicate peaks centered on neighboring planes. Grey X-marks indicate signals that are not sequence specifically assigned.

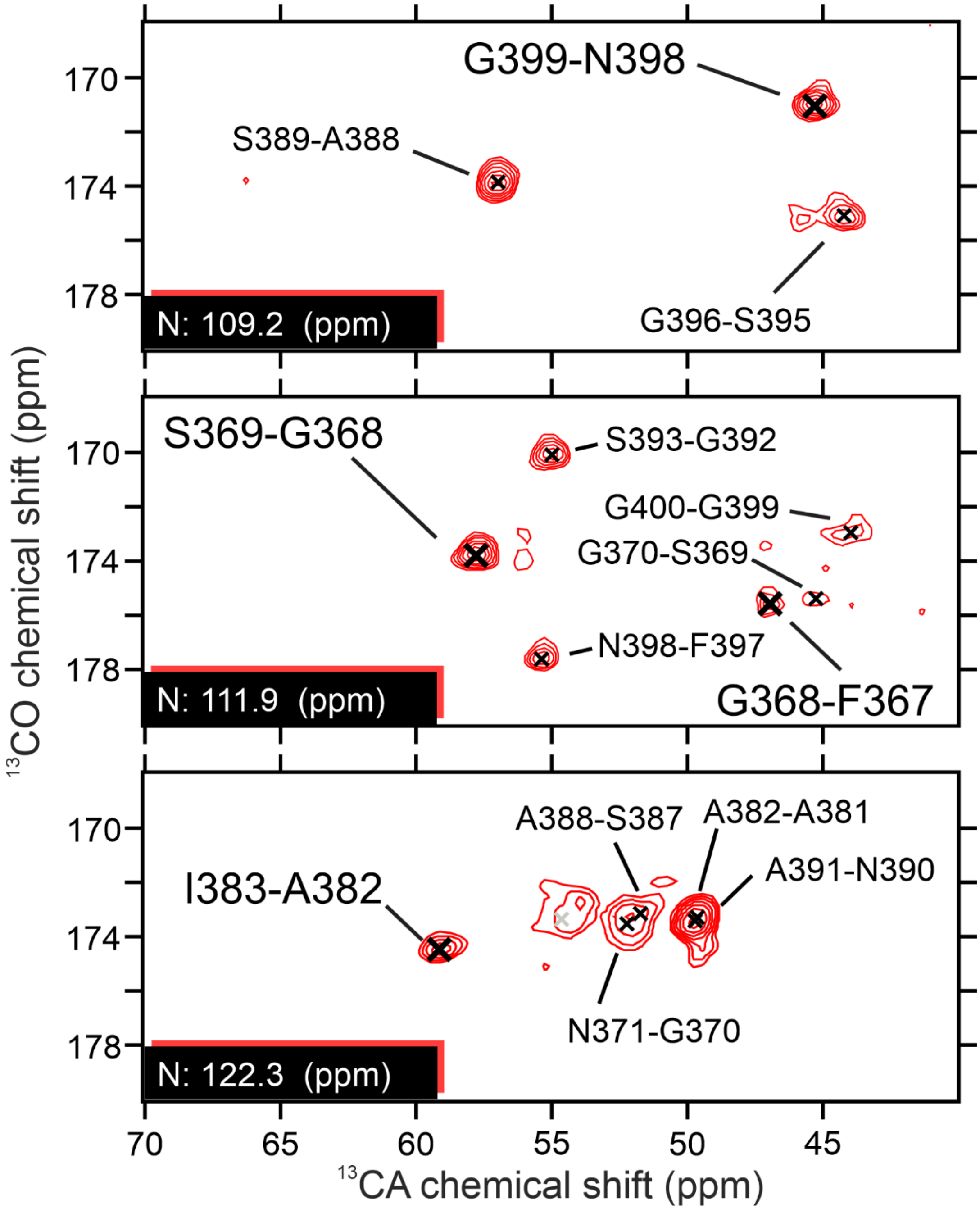

**Supplemental Figure 8. Solid state NMR chemical shift differences between TDP-43 fibril preparations.** Aggregation core corresponds to the observed chemical shifts for the residue 311–360 fragment of TDP-43-LC from reference [20]. *E. coli* TDP-43-LC corresponds to the observed chemical shifts from reference [42]. Flash Diluted TDP-43-LC corresponds to the observed chemical shifts from this paper. Lines connect the symbols in each plot to guide the eye between successive residues. Differences are only reported for residues that are structured in both samples in the comparison, although if one of the N, CO, CA, or CB atoms is missing from one or both data sets, a value of zero is reported and indicated with an asterisk. For example, the multiple differences of zero for the CB plot between residues 365–398 partly arise from the eleven Gly residues in this region that have no CB atom.

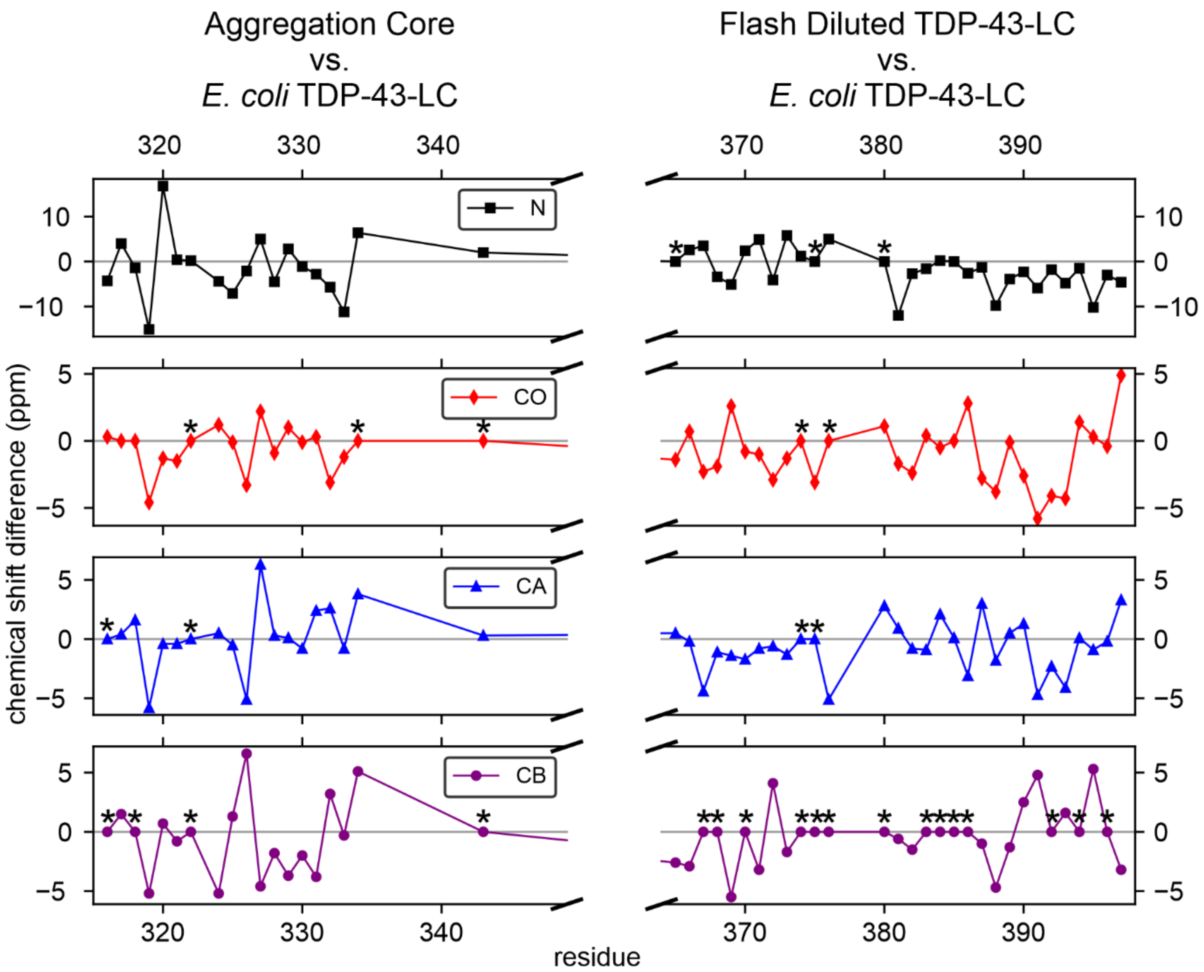

**Supplemental Table 1. Unambiguous resonance assignments for TDP-43-LC fibrils.**

| Residue | | $^{15}$N[a] | $^{13}$CA[a] | $^{13}$CO[a] | $^{13}$CB[a] | $^{13}$CG[a] | S/N[b] |
|---|---|---|---|---|---|---|---|
| Q | 365 | | 54.8 | 172.7 | 33.3 | | |
| A | 366 | 127.8 | 50.1 | 175.3 | 19.1 | | 8 |
| F | 367 | 124.6 | 51.2 | 175.6 | | | 12 |
| G | 368 | 112.0 | 47.1 | 173.6 | | | 8 |
| S | 369 | 111.9 | 57.8 | 175.6 | 63.0 | | 22 |
| G | 370 | 111.2 | 45.6 | 173.6 | | | 13 |
| N | 371 | 123.4 | 52.0 | 174.2 | 39.2 | 176.3 | 15 |
| N | 372 | 117.6 | 52.3 | 173.7 | 42.5 | 175.4 | 16 |
| S | 373 | 118.0 | 55.5 | 170.9 | 65.2 | | 22 |
| Y | 374 | 124.9 | 57.3 | 175.5 | 39.1 | | 10 |
| S | 375 | 120.9 | 55.6 | 172.6 | 64.5 | | 14 |
| G | 376 | 114.5 | 43.4 | | | | 8 |
| S | 377 | | | | | | |
| N | 378 | | | | | | |
| S | 379 | | | | | | |
| G | 380 | | 47.4 | 172.0 | | | |
| A | 381 | 120.6 | 51.8 | 173.2 | 19.1 | | 25 |
| A | 382 | 123.2 | 49.6 | 174.5 | 20.4 | | 29[c] |
| I | 383 | 122.4 | 59.2 | 174.8 | | | 13 |
| G | 384 | 116.7 | 46.7 | 173.4 | | | 10 |
| W | 385 | 126.3 | 59.2 | 174.8 | | | 9 |
| G | 386 | 104.4 | 44.4 | 175.8 | | | 8 |
| S | 387 | 119.3 | 58.6 | 173.1 | 63.0 | | 13 |
| A | 388 | 121.5 | 52.2 | 173.8 | 16.0 | | 23 |
| S | 389 | 109.5 | 57.1 | 173.3 | 66.6 | | 21 |
| N | 390 | 118.0 | 52.5 | 173.5 | 42.7 | 175.2 | 24 |
| A | 391 | 122.8 | 49.8 | 175.0 | 23.0 | | 28[c] |
| G | 392 | 107.2 | 44.3 | 170.0 | | | 17 |
| S | 393 | 111.5 | 55.1 | 171.3 | 65.8 | | 21 |
| G | 394 | 107.5 | 44.7 | 173.3 | | | 12 |
| S | 395 | 112.9 | 56.0 | 175.2 | 64.3 | | 12 |
| G | 396 | 109.0 | 44.3 | 174.2 | | | 15 |
| F | 397 | 113.1 | 57.3 | 177.5 | 35.7 | | 9 |
| N | 398 | 112.1 | 55.4 | 171.1 | 40.5 | 176.6 | 12 |
| G | 399 | 109.4 | 45.3 | 172.9 | | | 19 |
| G | 400 | 111.4 | 44.2 | 172.8 | | | 10 |
| F | 401 | 128.5 | | | | | |

[a] Chemical shift values are the average of the observed shifts from the NCACX, NCOCX, and CANCO 3D spectra.
[b] Signal to noise (S/N) values are values reported from Sparky for the 3D CANCO spectrum.
[c] These peaks arise from a highly overlapped site in the CANCO spectrum and therefore have an inflated S/N figure.

**Supplemental Table 2. Unassigned signals from the 3D CP-NCACX spectrum of TDP-43-LC fibrils**.

| $^{15}N$ | $^{13}CA$ | $^{13}CO$ | $^{13}CB$ | S/N[a] | Residue Type |
|---|---|---|---|---|---|
| 111.4 | 44.1 | 171.5 | | 8 | G |
| 113.9 | 55.5 | 173.0 | 65.4 | 10 | S |
| 114.9 | 59.1 | 177.7 | | 12 | RNDQEHILKMFPWYS |
| 114.9 | 46.2 | 173.0 | | 9 | G |
| 115.2 | 55.4 | 173.2 | 64.3 | 9 | S |
| 117.7 | 56.4 | 171.8 | 67.0 | 7 | S |
| 118.6 | 53.6 | 173.4 | 37.5 | 10 | RNDQEHILKMFPWY |
| 119.7 | 55.5 | 173.2 | 65.8 | 12 | S |
| 119.8 | 58.7 | 174.1 | 62.9 | 10 | S |
| 121.0 | 55.8 | 174.5 | | 11 | RNDQEHILKMFPWYS |
| 121.1 | 54.4 | 179.6 | 17.7 | 6 | A |

[a] Signal to noise (S/N) values are values reported from Sparky for the 3D CANCO spectrum.

**Supplemental Table 3. NCACX signal table for MCASSIGN.**

| Observed Chemical Shift (ppm) | | | | | Chemical Shift Uncertainty (ppm) | | | | | | |
|---|---|---|---|---|---|---|---|---|---|---|---|
| $^{15}N$ | $^{13}CA$ | $^{13}CO$ | $^{13}CB$ | $^{13}CG$ | $^{15}N$ | $^{13}CA$ | $^{13}CO$ | $^{13}CB$ | $^{13}CG$ | MP[a] | Residue Type |
| 104.3 | 44.2 | 175.7 | 1111.1 | 1111.1 | 0.4 | 0.3 | 0.3 | 0.3 | 0.3 | 1 | G |
| 107.0 | 44.2 | 170.0 | 1111.1 | 1111.1 | 0.4 | 0.3 | 0.3 | 0.3 | 0.3 | 1 | G |
| 107.4 | 44.6 | 173.5 | 1111.1 | 1111.1 | 0.5 | 0.3 | 0.6 | 0.3 | 0.3 | 1 | G |
| 108.9 | 44.2 | 174.2 | 1111.1 | 1111.1 | 0.4 | 0.5 | 0.5 | 0.3 | 0.3 | 1 | G |
| 109.5 | 45.3 | 172.9 | 1111.1 | 1111.1 | 0.4 | 0.3 | 0.3 | 0.3 | 0.3 | 1 | G |
| 109.4 | 57.0 | 173.4 | 66.5 | 1111.1 | 0.4 | 0.3 | 0.3 | 0.3 | 0.3 | 1 | S |
| 111.3 | 45.5 | 173.7 | 1111.1 | 1111.1 | 0.5 | 0.4 | 0.5 | 0.3 | 0.3 | 1 | G |
| 111.3 | 43.9 | 173.2 | 1111.1 | 1111.1 | 0.4 | 0.4 | 1.0 | 0.3 | 0.3 | 1 | G |
| 111.4 | 44.1 | 171.5 | 1111.1 | 1111.1 | 0.4 | 0.4 | 0.7 | 0.3 | 0.3 | 2 | G |
| 111.6 | 55.0 | 171.2 | 65.6 | 1111.1 | 0.4 | 0.3 | 0.5 | 0.3 | 0.3 | 1 | S |
| 111.9 | 46.9 | 173.6 | 1111.1 | 1111.1 | 0.4 | 0.5 | 0.5 | 0.3 | 0.3 | 1 | G |
| 111.9 | 57.7 | 175.5 | 62.8 | 1111.1 | 0.4 | 0.3 | 0.5 | 0.3 | 0.3 | 1 | S |
| 112.1 | 55.4 | 171.2 | 40.4 | 176.5 | 0.4 | 0.3 | 0.5 | 0.3 | 0.3 | 1 | ND |
| 112.9 | 56.0 | 175.2 | 64.3 | 1111.1 | 0.4 | 0.3 | 0.4 | 0.3 | 0.3 | 1 | S |
| 113.2 | 57.3 | 177.6 | 35.7 | 1111.1 | 0.4 | 0.3 | 0.5 | 0.5 | 0.3 | 1 | RNDQEHILKMFPWY |
| 113.9 | 55.5 | 173.0 | 65.4 | 1111.1 | 0.4 | 0.4 | 1.0 | 0.4 | 0.3 | 1 | S |
| 114.9 | 59.1 | 177.7 | 1111.1 | 1111.1 | 0.4 | 0.3 | 0.3 | 0.3 | 0.3 | 1 | RNDQEHILKMFPWYS |
| 114.9 | 46.2 | 173.0 | 1111.1 | 1111.1 | 0.4 | 0.3 | 0.4 | 0.3 | 0.3 | 1 | G |
| 115.2 | 55.4 | 173.2 | 64.3 | 1111.1 | 0.5 | 0.3 | 1.0 | 0.3 | 0.3 | 1 | S |
| 116.7 | 46.5 | 173.3 | 1111.1 | 1111.1 | 0.4 | 0.3 | 0.4 | 0.3 | 0.3 | 1 | G |
| 117.6 | 52.3 | 173.6 | 42.4 | 175.3 | 0.6 | 0.3 | 0.4 | 0.5 | 0.4 | 2 | ND |
| 117.7 | 56.4 | 171.8 | 67.0 | 1111.1 | 0.4 | 0.3 | 0.5 | 0.3 | 0.3 | 1 | S |
| 117.9 | 55.5 | 170.7 | 65.0 | 1111.1 | 0.4 | 0.3 | 0.3 | 0.4 | 0.3 | 1 | S |
| 118.6 | 53.6 | 173.4 | 37.5 | 1111.1 | 0.4 | 0.3 | 0.7 | 0.3 | 0.3 | 1 | RNDQEHILKMFPWY |
| 119.3 | 58.5 | 173.2 | 62.8 | 1111.1 | 0.5 | 0.4 | 0.5 | 0.4 | 0.3 | 1 | S |
| 119.7 | 55.5 | 173.2 | 65.8 | 1111.1 | 0.4 | 0.4 | 0.7 | 0.3 | 0.3 | 1 | S |
| 119.8 | 58.7 | 174.1 | 62.9 | 1111.1 | 0.5 | 0.4 | 0.5 | 0.4 | 0.3 | 1 | S |
| 120.5 | 51.7 | 173.1 | 19.1 | 1111.1 | 0.4 | 0.5 | 0.5 | 0.5 | 0.3 | 1 | A |
| 120.8 | 55.5 | 172.6 | 64.4 | 1111.1 | 0.4 | 0.3 | 0.5 | 0.4 | 0.3 | 1 | S |
| 121.0 | 55.8 | 174.5 | 1111.1 | 1111.1 | 0.4 | 0.3 | 0.5 | 0.4 | 0.3 | 1 | RNDQEHILKMFPWYS |
| 121.1 | 54.4 | 179.6 | 17.7 | 1111.1 | 0.4 | 0.3 | 0.4 | 0.3 | 0.3 | 1 | A |
| 121.5 | 52.1 | 173.7 | 16.0 | 1111.1 | 0.4 | 0.4 | 0.6 | 0.4 | 0.3 | 1 | A |
| 122.5 | 59.2 | 174.7 | 1111.1 | 1111.1 | 0.4 | 0.3 | 0.3 | 0.3 | 0.3 | 1 | RNDQEHILKMFPWYS |
| 122.6 | 49.8 | 174.9 | 22.9 | 1111.1 | 0.4 | 0.3 | 0.6 | 0.3 | 0.3 | 1 | A |
| 123.4 | 51.9 | 174.1 | 39.1 | 176.2 | 0.4 | 0.4 | 0.7 | 0.3 | 0.5 | 1 | ND |
| 123.2 | 49.4 | 174.6 | 20.3 | 1111.1 | 0.4 | 0.4 | 0.5 | 0.3 | 0.3 | 1 | A |
| 124.5 | 51.1 | 175.5 | 1111.1 | 1111.1 | 0.4 | 0.3 | 0.5 | 0.3 | 0.3 | 1 | RNDQEHILKMFPWYS |
| 125.0 | 57.1 | 175.5 | 1111.1 | 1111.1 | 0.5 | 0.4 | 0.4 | 0.3 | 0.3 | 1 | RNDQEHILKMFPWYS |
| 126.4 | 59.2 | 174.8 | 1111.1 | 1111.1 | 0.5 | 0.4 | 0.4 | 0.3 | 0.3 | 1 | RNDQEHILKMFPWYS |
| 127.9 | 50.1 | 175.2 | 19.0 | 1111.1 | 0.5 | 0.3 | 0.4 | 0.3 | 0.3 | 1 | A |

[a] multiplicity (MP), the number of times a signal can be assigned

**Supplemental Table 4. NCOCX signal table for MCASSIGN.**

| Observed Chemical Shift (ppm) | | | | | Chemical Shift Uncertainty (ppm) | | | | | | |
|---|---|---|---|---|---|---|---|---|---|---|---|
| $^{15}N$ | $^{13}CA$ | $^{13}CO$ | $^{13}CB$ | $^{13}CG$ | $^{15}N$ | $^{13}CA$ | $^{13}CO$ | $^{13}CB$ | $^{13}CG$ | MP[a] | |
| 107.1 | 50.0 | 175.1 | 23.0 | 1111.1 | 0.4 | 0.3 | 0.4 | 0.3 | 0.3 | 1 | A |
| 107.4 | 55.3 | 171.4 | 65.9 | 1111.1 | 0.4 | 0.3 | 0.3 | 0.3 | 0.3 | 1 | S |
| 109.1 | 56.1 | 175.2 | 64.3 | 1111.1 | 0.4 | 0.3 | 0.4 | 0.3 | 0.3 | 1 | S |
| 109.4 | 55.5 | 171.0 | 40.5 | 176.6 | 0.4 | 0.3 | 0.4 | 0.4 | 0.5 | 1 | ND |
| 109.5 | 52.3 | 173.8 | 16.0 | 1111.1 | 0.4 | 0.4 | 0.4 | 0.4 | 0.3 | 1 | A |
| 111.2 | 57.8 | 175.7 | 63.1 | 1111.1 | 0.4 | 0.3 | 0.4 | 0.3 | 0.3 | 1 | S |
| 111.3 | 44.4 | 170.1 | 1111.1 | 1111.1 | 0.4 | 0.3 | 0.4 | 0.3 | 0.3 | 1 | G |
| 111.6 | 45.4 | 172.8 | 1111.1 | 1111.1 | 0.4 | 0.5 | 0.4 | 0.3 | 0.3 | 1 | G |
| 112.0 | 47.3 | 173.5 | 1111.1 | 1111.1 | 0.4 | 0.3 | 0.4 | 0.3 | 0.3 | 1 | G |
| 112.1 | 51.5 | 175.7 | 1111.1 | 1111.1 | 0.4 | 0.3 | 0.5 | 0.3 | 0.3 | 1 | RNDQEHILKMFPWYS |
| 112.1 | 57.4 | 177.4 | 1111.1 | 1111.1 | 0.4 | 0.4 | 0.4 | 0.3 | 0.3 | 1 | RNDQEHILKMFPWYS |
| 112.9 | 44.6 | 173.3 | 1111.1 | 1111.1 | 0.4 | 0.7 | 1.0 | 0.3 | 0.3 | 1 | G |
| 113.1 | 44.6 | 173.9 | 1111.1 | 1111.1 | 0.4 | 0.7 | 1.0 | 0.3 | 0.3 | 1 | G |
| 114.3 | 55.8 | 172.6 | 64.5 | 1111.1 | 0.4 | 0.3 | 0.5 | 0.3 | 0.3 | 1 | S |
| 114.5 | 56.9 | 171.5 | 67.2 | 1111.1 | 0.4 | 0.3 | 0.4 | 0.3 | 0.3 | 1 | S |
| 114.9 | 57.6 | 172.7 | 1111.1 | 1111.1 | 0.5 | 0.3 | 0.4 | 0.3 | 0.3 | 1 | RNDQEHILKMFPWYS |
| 115.0 | 53.9 | 173.1 | 1111.1 | 1111.1 | 0.4 | 0.4 | 0.4 | 0.3 | 0.3 | 1 | RNDQEHILKMFPWYS |
| 117.0 | 59.2 | 174.8 | 1111.1 | 1111.1 | 0.4 | 0.4 | 0.4 | 0.4 | 0.3 | 1 | RNDQEHILKMFPWYS |
| 117.5 | 52.2 | 174.3 | 39.2 | 176.4 | 0.4 | 0.4 | 0.3 | 0.3 | 0.3 | 1 | ND |
| 118.0 | 46.6 | 173.3 | 1111.1 | 1111.1 | 0.4 | 0.4 | 0.4 | 0.3 | 0.3 | 1 | G |
| 118.1 | 57.1 | 173.1 | 66.6 | 1111.1 | 0.4 | 0.3 | 0.4 | 0.3 | 0.3 | 1 | S |
| 118.2 | 52.4 | 173.7 | 42.5 | 175.5 | 0.4 | 0.3 | 0.5 | 0.4 | 0.3 | 1 | ND |
| 118.1 | 55.6 | 173.7 | 42.5 | 1111.1 | 0.4 | 0.3 | 0.5 | 0.4 | 0.3 | 1 | RNDQEHILKMFPWY |
| 119.3 | 59.2 | 174.7 | 62.7 | 1111.1 | 0.4 | 0.3 | 0.5 | 0.3 | 0.3 | 1 | S |
| 119.5 | 44.6 | 175.9 | 1111.1 | 1111.1 | 0.4 | 0.3 | 0.5 | 0.3 | 0.3 | 1 | G |
| 120.7 | 47.4 | 171.9 | 1111.1 | 1111.1 | 0.4 | 0.3 | 0.4 | 0.3 | 0.3 | 1 | G |
| 120.9 | 57.5 | 175.5 | 39.1 | 1111.1 | 0.4 | 0.3 | 0.5 | 0.3 | 0.3 | 1 | RNDQEHILKMFPWY |
| 121.4 | 54.5 | 179.8 | 17.6 | 1111.1 | 0.4 | 0.3 | 0.3 | 0.3 | 0.3 | 1 | A |
| 121.5 | 54.5 | 179.8 | 17.5 | 1111.1 | 0.4 | 0.3 | 0.4 | 0.3 | 0.3 | 1 | A |
| 121.6 | 58.6 | 173.2 | 63.1 | 1111.1 | 0.4 | 0.3 | 0.5 | 0.3 | 0.3 | 1 | S |
| 122.4 | 49.8 | 174.5 | 20.4 | 1111.1 | 0.4 | 0.3 | 0.4 | 0.3 | 0.3 | 1 | A |
| 122.6 | 52.6 | 173.6 | 42.9 | 175.0 | 0.4 | 0.3 | 0.4 | 0.3 | 0.5 | 1 | ND |
| 123.3 | 52.0 | 173.4 | 19.1 | 1111.1 | 0.4 | 0.3 | 0.5 | 0.3 | 0.3 | 1 | A |
| 123.4 | 45.7 | 173.6 | 1111.1 | 1111.1 | 0.4 | 0.3 | 0.4 | 0.3 | 0.3 | 1 | G |
| 124.7 | 50.3 | 175.4 | 19.2 | 1111.1 | 0.4 | 0.3 | 0.4 | 0.3 | 0.3 | 1 | A |
| 124.9 | 55.5 | 171.1 | 65.3 | 1111.1 | 0.4 | 0.3 | 0.3 | 0.3 | 0.3 | 1 | S |
| 126.0 | 46.8 | 173.4 | 1111.1 | 1111.1 | 0.4 | 0.3 | 0.3 | 0.3 | 0.3 | 1 | G |
| 127.7 | 54.8 | 172.8 | 33.3 | 1111.1 | 0.4 | 0.5 | 0.5 | 0.5 | 0.3 | 1 | RNDQEHILKMFPWY |
| 128.5 | 44.4 | 172.4 | 1111.1 | 1111.1 | 0.4 | 0.4 | 0.4 | 0.3 | 0.3 | 1 | G |

[a] multiplicity (MP), the number of times a signal can be assigned

**Supplemental Table 5. CANCO signal table for MCASSIGN.**

| Observed Chemical Shift (ppm) | | | Chemical Shift Uncertainty (ppm) | | | | |
|---|---|---|---|---|---|---|---|
| $^{15}N$ | $^{13}CA$ | $^{13}CO$ | $^{15}N$ | $^{13}CA$ | $^{13}CO$ | MP[a] | |
| 104.5 | 44.4 | 174.8 | 0.4 | 0.4 | 0.4 | 1 | G |
| 107.4 | 43.3 | 171.3 | 0.4 | 0.4 | 0.4 | 1 | G |
| 107.4 | 44.3 | 175.0 | 0.4 | 0.4 | 0.4 | 2 | G |
| 107.7 | 44.9 | 171.4 | 0.4 | 0.4 | 0.4 | 1 | G |
| 109.1 | 44.2 | 175.2 | 0.4 | 0.4 | 0.4 | 1 | G |
| 109.3 | 45.3 | 171.0 | 0.4 | 0.4 | 0.4 | 1 | G |
| 109.5 | 57.1 | 173.8 | 0.4 | 0.4 | 0.4 | 1 | S |
| 111.2 | 45.7 | 175.5 | 0.4 | 0.4 | 0.4 | 1 | G |
| 110.8 | 44.4 | 175.0 | 0.4 | 0.4 | 0.4 | 1 | G |
| 111.4 | 44.2 | 172.9 | 0.4 | 0.4 | 0.4 | 1 | G |
| 111.6 | 55.1 | 170.0 | 0.4 | 0.4 | 0.4 | 1 | S |
| 111.9 | 47.0 | 175.6 | 0.4 | 0.4 | 0.4 | 1 | G |
| 111.9 | 57.8 | 173.8 | 0.4 | 0.4 | 0.4 | 1 | S |
| 112.2 | 55.4 | 177.6 | 0.4 | 0.4 | 0.4 | 1 | ND |
| 112.9 | 56.0 | 173.1 | 0.4 | 0.4 | 1.0 | 1 | S |
| 113.1 | 57.3 | 174.4 | 0.4 | 0.4 | 0.4 | 1 | RNDQEHILKMFPWYS |
| 114.5 | 56.1 | 171.8 | 0.6 | 0.6 | 1.0 | 2 | S |
| 114.7 | 43.4 | 172.7 | 0.4 | 0.4 | 0.4 | 1 | G |
| 114.9 | 59.2 | 173.1 | 0.4 | 0.4 | 0.4 | 1 | RNDQEHILKMFPWYS |
| 115.0 | 46.2 | 172.7 | 0.4 | 0.4 | 0.4 | 1 | G |
| 115.4 | 54.9 | 173.8 | 0.4 | 0.4 | 0.4 | 1 | S |
| 116.5 | 46.8 | 174.9 | 0.4 | 0.4 | 0.4 | 1 | G |
| 117.2 | 56.4 | 172.1 | 0.4 | 0.4 | 0.4 | 1 | S |
| 117.6 | 52.2 | 174.3 | 0.4 | 0.4 | 0.4 | 1 | ND |
| 118.2 | 52.5 | 173.4 | 0.4 | 0.4 | 0.4 | 1 | ND |
| 118.0 | 55.4 | 173.7 | 0.4 | 0.4 | 0.4 | 1 | S |
| 119.2 | 58.6 | 175.8 | 0.5 | 0.4 | 0.4 | 1 | S |
| 119.1 | 53.6 | 174.4 | 0.4 | 0.4 | 0.4 | 1 | RNDQEHILKMFPWYS |
| 119.8 | 55.7 | 173.4 | 0.4 | 0.4 | 0.4 | 1 | S |
| 120.2 | 58.9 | 174.7 | 0.5 | 0.4 | 0.4 | 1 | S |
| 120.7 | 51.7 | 172.0 | 0.4 | 0.4 | 0.4 | 1 | A |
| 120.9 | 55.6 | 175.6 | 0.4 | 0.4 | 0.4 | 1 | S |
| 121.1 | 54.6 | 179.8 | 0.4 | 0.4 | 0.4 | 1 | A |
| 121.3 | 55.4 | 173.2 | 0.4 | 0.4 | 0.4 | 1 | RNDQEHILKMFPWYS |
| 121.4 | 52.1 | 173.0 | 0.6 | 0.4 | 0.6 | 2 | A |
| 122.4 | 59.1 | 174.5 | 0.4 | 0.4 | 0.4 | 1 | RNDQEHILKMFPWYS |
| 123.1 | 49.6 | 173.2 | 0.6 | 0.5 | 0.5 | 2 | A |
| 123.3 | 52.0 | 173.6 | 0.4 | 0.4 | 0.4 | 1 | ND |
| 124.5 | 51.1 | 175.3 | 0.4 | 0.4 | 0.4 | 1 | RNDQEHILKMFPWYS |
| 124.8 | 57.3 | 170.8 | 0.4 | 0.4 | 0.4 | 1 | RNDQEHILKMFPWYS |
| 126.5 | 59.2 | 173.4 | 0.4 | 0.4 | 0.4 | 1 | RNDQEHILKMFPWYS |
| 127.8 | 49.8 | 172.5 | 0.4 | 0.4 | 0.4 | 1 | A |

[a] multiplicity (MP), the number of times a signal can be assigned

**Supplemental Table S6. Solid state NMR data collection and processing parameters.**

| | Spectrum | Acquisition Parameters[a,b] | Processing Parameters[c] |
|---|---|---|---|
| **Aged Droplet** | 2D $^{13}$C-$^{13}$C CP-DARR | ns=112; $\tau_{aq}$=10.24 ms; $\nu_{1H\text{-}carr}$=2.5 ppm; $\nu_{13C\text{-}carr}$=99.0 ppm; $\nu_{1H\text{-}CP}$=63 kHz; $\nu_{13C\text{-}CP}$=50 kHz; $\nu_{1Hdec}$=83.3 kHz; $\nu_{DARR}$=13 kHz; $\tau_{1H\text{-}\pi/2}$=3 µs; $\tau_{13C\text{-}\pi/2}$=4 µs; $\tau_{CP}$=1.0 ms; $\tau_{DARR}$=50 ms; $\tau_{1H\text{-}dec}$=6.0 µs; $\Delta t_1$=24.0 µs; $\tau_{t1}$=5.4 ms; | $GLB_{t1}$=100 Hz; $GLB_{t2}$=100 Hz; LP in $t_2$ |
| | 2D CP-NCACX | ns=432; $\tau_{aq}$=10.24 ms; $\nu_{1H\text{-}carr}$=2.5 ppm; $\nu_{13C\text{-}carr}$=64.8 ppm; $\nu_{15N\text{-}carr}$=119.9 ppm; $\nu_{1H\text{-}CP}$=49 kHz; $\nu_{15N\text{-}CP}$=36 kHz; $\nu_{15N\text{-}SCP}$=36 kHz; $\nu_{13CA\text{-}SCP}$=23 kHz; $\nu_{1H\text{-}dec}$=83.3 kHz; $\nu_{DARR}$=13 kHz; $\tau_{1H\text{-}\pi/2}$=3 µs; $\tau_{13C\text{-}\pi/2}$=4 µs; $\tau_{15N\text{-}\pi/2}$=6 µs; $\tau_{CP}$=0.75 ms; $\tau_{SCP}$=4.0 ms; $\tau_{DARR}$=50 ms; $\tau_{1H\text{-}dec}$=6.0 µs; $\Delta t_1$=108.0 µs; $\tau_{t1}$=11.0 ms; | $GLB_{t1}$=60 Hz; $GLB_{t2}$=100 Hz; |
| | 2D CP-NCOCX | ns=432; $\tau_{aq}$=10.24 ms; $\nu_{1H\text{-}carr}$=2.5 ppm; $\nu_{13C\text{-}carr}$=180.4 ppm; $\nu_{15N\text{-}carr}$=119.9 ppm; $\nu_{1H\text{-}CP}$=49 kHz; $\nu_{15N\text{-}CP}$=36 kHz; $\nu_{15N\text{-}SCP}$=37 kHz; $\nu_{13CO\text{-}SCP}$=50 kHz; $\nu_{1H\text{-}dec}$=83.3 kHz; $\nu_{DARR}$=13 kHz; $\tau_{1H\text{-}\pi/2}$=3 µs; $\tau_{13C\text{-}\pi/2}$=4 µs; $\tau_{15N\text{-}\pi/2}$=6 µs; $\tau_{CP}$=0.75 ms; $\tau_{SCP}$=4.0 ms; $\tau_{DARR}$=50 ms; $\tau_{1H\text{-}dec}$=6.0 µs; $\Delta t_1$=108.0 µs; $\tau_{t1}$=11.0 ms; | $GLB_{t1}$=60 Hz; $GLB_{t2}$=100 Hz; |
| | 2D $^{1}$H-$^{13}$C INEPT | ns=32; $\tau_{aq}$=30.72 ms; $\nu_{1H\text{-}carr}$=2.5 ppm; $\nu_{13C\text{-}carr}$=95.3 ppm; $\nu_{1H\text{-}dec}$=10.0 kHz; $\tau_{1H\text{-}\pi/2}$=3 µs; $\tau_{13C\text{-}\pi/2}$=4 µs; $\tau_{J/2}$=0.76 ms; $\Delta t_1$=50.0 µs; $\tau_{t1}$=11.25 ms; | $GLB_{t1}$=10 Hz; $GLB_{t2}$=50 Hz; |
| **Flash Diluted** | 2D $^{13}$C-$^{13}$C CP-DARR | ns=96; $\tau_{aq}$=10.24 ms; $\nu_{1H\text{-}carr}$=4.4 ppm; $\nu_{13C\text{-}carr}$=94.6 ppm; $\nu_{1H\text{-}CP}$=63 kHz; $\nu_{13C\text{-}CP}$=50 kHz; $\nu_{1Hdec}$=83.3 kHz; $\nu_{DARR}$=13 kHz; $\tau_{1H\text{-}\pi/2}$=3 µs; $\tau_{13C\text{-}\pi/2}$=4 µs; $\tau_{CP}$=1.0 ms; $\tau_{DARR}$=50 ms; $\tau_{1H\text{-}dec}$=5.7 µs; $\Delta t_1$=22.8 µs; $\tau_{t1}$=6.0 ms; | $GLB_{t1}$=100 Hz; $GLB_{t2}$=100 Hz; LP in $t_2$ |
| | 2D CP-NCACX | ns=384; $\tau_{aq}$=10.24 ms; $\nu_{1H\text{-}carr}$=4.4 ppm; $\nu_{13C\text{-}carr}$=56.1 ppm; $\nu_{15N\text{-}carr}$=117.4 ppm; $\nu_{1H\text{-}CP}$=49 kHz; $\nu_{15N\text{-}CP}$=36 kHz; $\nu_{15N\text{-}SCP}$=34 kHz; $\nu_{13CA\text{-}SCP}$=21 kHz; $\nu_{1H\text{-}dec}$=83.3 kHz; $\nu_{DARR}$=13 kHz; $\tau_{1H\text{-}\pi/2}$=3 µs; $\tau_{13C\text{-}\pi/2}$=4 µs; $\tau_{15N\text{-}\pi/2}$=6 µs; $\tau_{CP}$=0.75 ms; $\tau_{SCP}$=4.0 ms; $\tau_{DARR}$=50 ms; $\tau_{1H\text{-}dec}$=5.7 µs; $\Delta t_1$=114.0 µs; $\tau_{t1}$=11.1 ms; | $GLB_{t1}$=60 Hz; $GLB_{t2}$=100 Hz; |
| | 2D CP-NCOCX | ns=416; $\tau_{aq}$=10.24 ms; $\nu_{1H\text{-}carr}$=4.4 ppm; $\nu_{13C\text{-}carr}$=175.6 ppm; $\nu_{15N\text{-}carr}$=117.4 ppm; $\nu_{1H\text{-}CP}$=49 kHz; $\nu_{15N\text{-}CP}$=36 kHz; $\nu_{15N\text{-}SCP}$=38 kHz; $\nu_{13CO\text{-}SCP}$=51 kHz; $\nu_{1H\text{-}dec}$=83.3 kHz; $\nu_{DARR}$=13 kHz; $\tau_{1H\text{-}\pi/2}$=3 µs; $\tau_{13C\text{-}\pi/2}$=4 µs; $\tau_{15N\text{-}\pi/2}$=6 µs; $\tau_{CP}$=0.75 ms; $\tau_{SCP}$=4.0 ms; $\tau_{DARR}$=50 ms; $\tau_{1H\text{-}dec}$=5.7 µs; $\Delta t_1$=114.0 µs; $\tau_{t1}$=11.1 ms; | $GLB_{t1}$=60 Hz; $GLB_{t2}$=100 Hz; LP in $t_2$ |

| | | | |
|---|---|---|---|
| | 2D $^{1}H$-$^{13}C$ INEPT | ns=64; $\tau_{aq}$=20.48 ms; $\nu_{1H-carr}$=6.2 ppm; $\nu_{13C-carr}$=96.5 ppm; $\nu_{1H-dec}$=12.5 kHz; $\tau_{1H-\pi/2}$=3 µs; $\tau_{13C-\pi/2}$=4 µs; $\tau_{J/2}$=1.0 ms; $\Delta t_1$=50.0 µs; $\tau_{t1}$=11.25 ms; | $GLB_{t1}$=10 Hz; $GLB_{t2}$=50 Hz; |
| | 3D CP-NCACX | ns=40; $\tau_{aq}$=10.24 ms; $\nu_{1H-carr}$=3.2 ppm; $\nu_{13C-carr}$=65.5 ppm; $\nu_{15N-carr}$=120.6 ppm; $\nu_{1H-CP}$=49 kHz; $\nu_{15N-CP}$=36 kHz; $\nu_{15N-SCP}$=35 kHz; $\nu_{13CA-SCP}$=22 kHz; $\nu_{1H-dec}$=83.3 kHz; $\nu_{DARR}$=13 kHz; $\tau_{1H-\pi/2}$=3 µs; $\tau_{13C-\pi/2}$=4 µs; $\tau_{15N-\pi/2}$=6 µs; $\tau_{CP}$=0.75 ms; $\tau_{SCP}$=4.0 ms; $\tau_{DARR}$=50 ms; $\tau_{1H-dec}$=6.0 µs; $\Delta t_2$=132.0 µs; $\tau_{t2}$=3.3 ms; $\Delta t_1$=180.0 µs; $\tau_{t1}$=7.6 ms; | $GLB_{t1}$=60 Hz; $GLB_{t2}$=100 Hz; $GLB_{t3}$=100 Hz; |
| | 3D CP-NCOCX | ns=48; $\tau_{aq}$=10.24 ms; $\nu_{1H-carr}$=3.2 ppm; $\nu_{13C-carr}$=176.0 ppm; $\nu_{15N-carr}$=120.6 ppm; $\nu_{1H-CP}$=49 kHz; $\nu_{15N-CP}$=36 kHz; $\nu_{15N-SCP}$=36 kHz; $\nu_{13CO-SCP}$=49 kHz; $\nu_{1H-dec}$=83.3 kHz; $\nu_{DARR}$=13 kHz; $\tau_{1H-\pi/2}$=3 µs; $\tau_{13C-\pi/2}$=4 µs; $\tau_{15N-\pi/2}$=6 µs; $\tau_{CP}$=0.75 ms; $\tau_{SCP}$=4.0 ms; $\tau_{DARR}$=50 ms; $\tau_{1H-dec}$=6.0 µs; $\Delta t_2$=132.0 µs; $\tau_{t2}$=3.8 ms; $\Delta t_1$=180.0 µs; $\tau_{t1}$=8.1 ms; | $GLB_{t1}$=100 Hz; $GLB_{t2}$=60 Hz; $GLB_{t3}$=100 Hz; |
| | 3D CP-CANCO | ns=16; $\tau_{aq}$=10.24 ms; $\nu_{1H-carr}$=3.0 ppm; $\nu_{13C-carr}$=65.3 ppm; $\nu_{15N-carr}$=120.4 ppm; $\nu_{1H-CP}$=63 kHz; $\nu_{13C-CP}$=50 kHz; $\nu_{15N-SCP}$=36 kHz; $\nu_{13CA-SCP}$=22 kHz; $\nu_{13CO-SCP-eff}$=49 kHz; $\nu_{1Hdec}$=83.3 kHz; $\tau_{1H-\pi/2}$=3 µs; $\tau_{SCP}$=4.0 ms; $\tau_{1H-dec}$=6.0 µs; $\Delta t_2$=180.0 µs; $\tau_{t2}$=8.1 ms; $\Delta t_1$=60.0 µs; $\tau_{t1}$=3.8 ms; | $GLB_{t1}$=100 Hz; $GLB_{t2}$=60 Hz; $GLB_{t3}$=100 Hz; |

[a] All $^{1}H$-X CP transfers used a 20% ramp on the $^{1}H$ channel. All $^{15}N$-$^{13}C$ SCP transfers used either a 5% or 10% ramp on the $^{15}N$ channel.

[b] ns, number of scans averaged; $\tau_{aq}$, total acquisition time in the direct dimension; $\nu_{1H-carr}$, $^{1}H$ carrier frequency; $\nu_{13C-carr}$, $^{13}C$ carrier frequency; $\nu_{15N-carr}$, $^{15}N$ carrier frequency; $\nu_{1H-CP}$, $^{1}H$ CP pulse power; $\nu_{13C-CP}$, $^{13}C$ CP pulse power; $\nu_{15N-CP}$, $^{15}N$ CP pulse power; $\nu_{15N-SCP}$, $^{15}N$ Specific-CP pulse power; $\nu_{13CA-SCP}$, $^{13}C$ α-carbon Specific-CP pulse power; $\nu_{13CO-SCP}$, $^{13}C$ carbonyl-carbon Specific-CP pulse power; $\nu_{1H-dec}$, $^{1}H$ decoupling pulse power; $\nu_{DARR}$, $^{13}C$-$^{13}C$ DARR $^{1}H$ pulse power; $\tau_{1H-\pi/2}$, $^{1}H$ π/2 pulse length; $\tau_{13C-\pi/2}$, $^{13}C$ π/2 pulse length; $\tau_{15N-\pi/2}$, $^{15}N$ π/2 pulse length; $\tau_{CP}$, $^{1}H$-X CP contact time; $\tau_{SCP}$, $^{15}N$-$^{13}C$ Specific-CP contact time; $\tau_{DARR}$, $^{13}C$-$^{13}C$ DARR mixing time; $\tau_{J/2}$,1/2 echo delay; $\tau_{1H-dec}$, $^{1}H$ SPINAL64 π pulse length; $\Delta t_2$, time increment in the $t_2$ dimension; $\tau_{t2}$, total acquisition time in the $t_2$ dimension; $\Delta t_1$, time increment in the $t_1$ dimension; $\tau_{t1}$, total acquisition time in the $t_1$ dimension;

[c] Zero filling was applied twice in each dimension prior to Fourier transformation. GLB, gaussian line broadening and LP, forward linear prediction with standard NMRPipe parameters.
